# Canonical Wnt signaling promotes formation of somatic permeability barrier for proper germ cell differentiation

**DOI:** 10.1101/2022.02.15.480616

**Authors:** Ting-An Chen, Kun-Yang Lin, Shun-Min Yang, Chen-Yuan Tseng, Yu-Ting Wang, Chi-Hung Lin, Lichao Luo, Yu Cai, Hwei-Jan Hsu

**Author notes:** Department of Biochemistry and Molecular Pharmacology, New York University Grossman School of Medicine, New York, NY 10016, USA. Haihe Biopharma Ltd., 421 Newton Road, Zhangjiang Hi-Tech Park, Pudong Shanghai, 201203, China (Lichao Luo). These authors contributed equally to this work (Ting-An Chen, Kun-Yang Lin, Shun-Min Yang). The order of names was determined alphabetically.

## Abstract

Morphogen-mediated signaling is critical for proper organ development and stem cell function, and well-characterized mechanisms spatiotemporally limit the expression of ligands, receptors, and ligand-binding cell-surface glypicans. Here, we show that in the developing *Drosophila* ovary, canonical Wnt signaling promotes the formation of somatic escort cells (ECs) and their protrusions, which establish a physical permeability barrier to define morphogen territories for proper germ cell differentiation. The protrusions shield germ cells from Dpp and Wingless morphogens produced by the germline stem cell (GSC) niche and normally only received by GSCs. Genetic disruption of EC protrusions allows GSC progeny to also receive Dpp and Wingless, which subsequently disrupt germ cell differentiation. Our results reveal a role for canonical Wnt signaling in specifying the ovarian somatic cells necessary for germ cell differentiation. Additionally, we demonstrate the morphogen-limiting function of this physical permeability barrier, which may be a common mechanism in other organs across species.

## Introduction

Morphogens are long-range signaling molecules that often establish gradients within developing tissues by traveling up to a few dozen cell diameters from the secreting cell source [1]. Along the gradient, cells receiving different concentrations of morphogen signal will display different target gene expression profiles and differentiate into distinct cell types. The establishment of such morphogen gradients depends largely on tightly regulated expression of morphogen receptors and negative regulators, though other mechanisms may participate as well. One important family of morphogens is the Wnt proteins, which control various aspects of tissue development and stem cell function in many adult tissues [2, 3]. However, relatively few studies have been conducted to describe the role of Wnt signaling in the development of ovaries, and the mechanisms controlling Wnt territory in the ovaries are unclear.

Wnt signaling is a highly conserved process that includes both canonical and non-canonical pathways [2, 4]. Canonical Wnt signaling is initiated by the binding of Wnt to Frizzled (Fz) receptors and Lipoprotein Receptor Protein (LRP) coreceptors. The Wnt-Fz-LRP complex recruits the scaffolding protein Dishevelled (Dsh in *Drosophila*), leading to disruption of the destruction complex and consequent stabilization of β-catenin (Armadillo, Arm, in *Drosophila*). The stable β-catenin protein translocates to the nucleus, where it forms a complex with the T-cell factor (TCF) transcription factor and co-activators, including Pygopus (Pygo), to regulate target gene expression. Wnts can also trigger a non-canonical, β-catenin-independent pathway by binding to a non-LRP coreceptor. Such non-canonical signaling can be further divided into the Planar Cell Polarity and the Wnt/Ca^2+^ pathways [5]. *Drosophila* has seven Wnt ligands with vertebrate orthologs: Wingless (Wg, ortholog of vertebrate Wnt1), Wnt2 (vertebrate Wnt7), Wnt3/5 (vertebrate Wnt5), Wnt4 (vertebrate Wnt9), Wnt6 (vertebrate Wnt6), Wnt8 (vertebrate Wnt8), and Wnt10 (vertebrate Wnt10) [6]. Several of these Wnt ligands have been shown to participate in primordial germ cell (PGC) development in different species. For example, Wnt3 functions in mouse PGC specification [7], while Wnt4 functions in female sex differentiation [8] and Wnt5 controls PGC migration [9]. The role of Wnt signaling in the adult germarium has been extensively studied [10–13]; however, fewer studies have examined the role of Wnt signaling during ovary development. In flies, Wg and Wnt8 respectively control embryonic PGC proliferation and migration [14, 15], whereas Wnt4 mediates non-canonical Wnt signaling to control the PGC-soma interaction in larval ovaries [16] and to stimulate apical cell migration during germarium formation in pupal ovaries [17]. Therefore, the role of canonical Wnt signaling in the development of ovaries is not fully understood.

We used the *Drosophila* ovary as a model to address the potential role and functional effects of canonical Wnt signaling in ovary development because of its simple cell composition and well-characterized cell biology (Fig. 1A) [18–20]. Thegonad of first instar larvae (L1; right after hatching) contains only a few PGCs and somatic gonad precursors (SGPs). Beginning at the early second instar larval stage (L2), a first morphogenesis occurs along the anterior-posterior and medial-lateral axes. A two-dimensional array of 16-20 stacks of somatic cells called terminal filaments (TFs) is generated by the end of third-instar larval (L3) stage, and the remaining SGPs differentiate into various somatic cell types. Apical cells are positioned above TFs. Intermingled cells (ICs) intermingle with PGCs, and basal cells are located at the bottom of the gonad. During pupal stages, apical cells migrate basally between TFs and through both ICs and basal cells to form 16-20 ovarioles [17]. Each ovariole bears 6-7 sequentially developing egg chambers and is considered to be a functional unit for producing eggs [21]. The anterior structure of the ovariole is called the germarium. Within this region, the ICs closest to the basal TFs differentiate into cap cells [22], whereas more distal ICs become escort cells (ECs). The cap cells, anterior-most ECs, and one TF constitute a GSC niche [23], which produces Decapentaplegic (Dpp, a *Drosophila* BMP), for the maintenance of two or three GSCs [24]. Each GSC is located directly adjacent to cap cells and contains a membranous organelle, called a fusome, which is found near the GSC-cap cell interface [25]. Each GSC division gives rise to one daughter GSC and one cystoblast (CB) that subsequently undergoes four rounds of incomplete division to become a 2-, 4-, 8-, and then 16-cell cyst [26] as it migrates through regions 1 and 2 of the germarium [27]. During these divisions, the fusome grows to interconnect the germ cells within the cyst, and it takes on a branched morphology in 4-, 8-and 16-cell cysts. Around the CB and cyst, ECs extend long cellular membrane protrusions to wrap and facilitate the differentiation of GSC progeny [28]. At the 2A/2B boundary, follicle cells substitute for ECs to wrap 16-cell cysts, which take on a lens-like shape. After acquiring a monolayer of follicle cells (derived from follicle stem cells at the 2A/2B boundary), the 16-cell cyst becomes a newly formed egg chamber and buds off from the germarium, eventually developing into a mature egg.

**Fig. 1.**
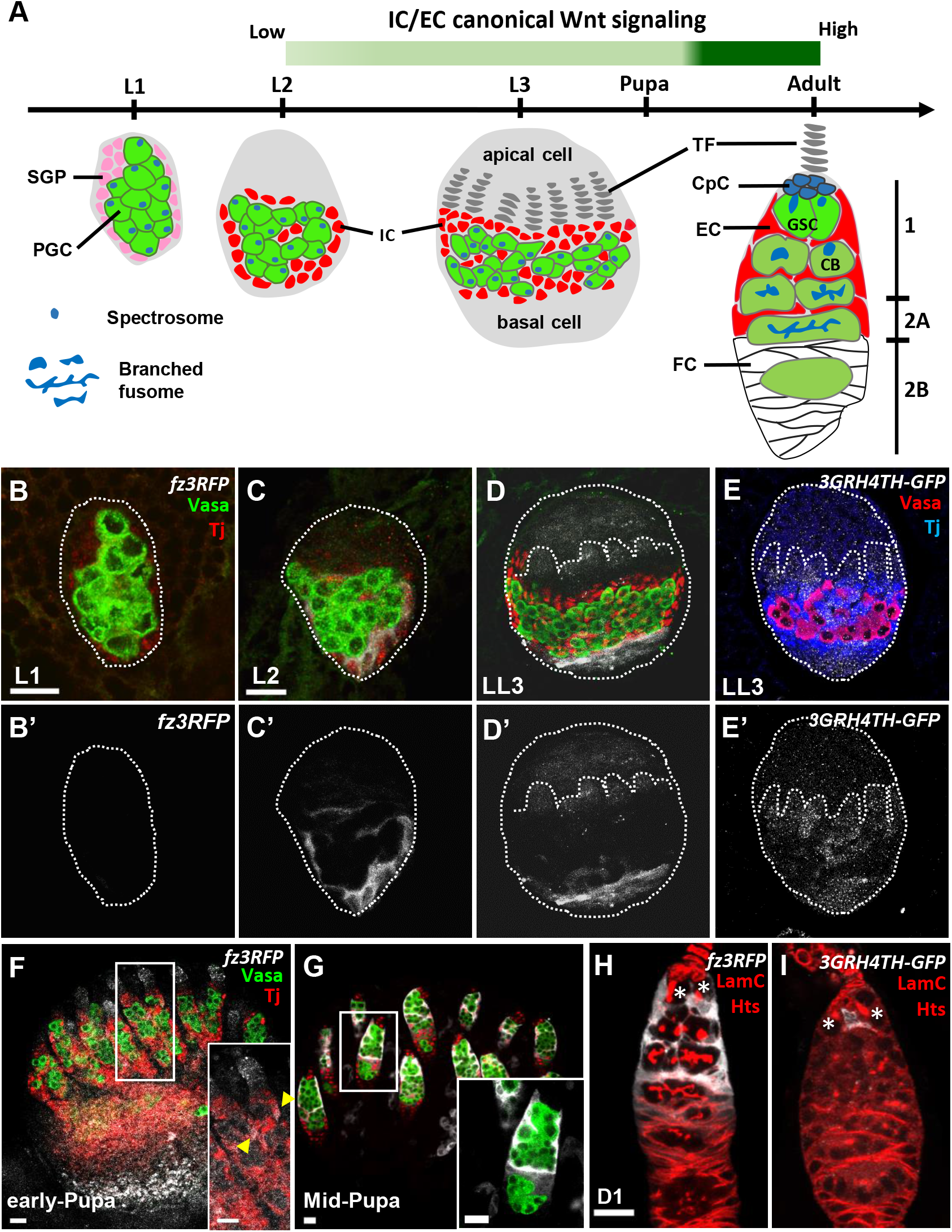
Canonical Wnt signaling is activated in the somatic cells of developing ovaries. **(A)** Schematic of *Drosophila* larval gonads and the adult germarium. Primordial germ cells (PGCs), each with a round-shaped fusome (unique membrane-enriched organelle), and somatic gonad precursors (SGPs) are present at the L1 and L2 stages. PGC and SGP numbers are greatly increased at the L3 stage, and SGPs differentiate into apical cells, terminal filaments (TFs), intermingled cells (ICs) and basal cells. ICs later further differentiate into cap cells (CpCs), escort cells (ECs), and probably follicle cells (FCs). During the pupal stage, apical cells migrate through TFs, ICs and basal cells to generate ovarioles; the anterior structure of the ovariole is the germarium. In the adult germarium, GSCs and their progeny are wrapped by EC protrusions in regions 1 and 2A. Then, ECs are replaced by follicle cells (FCs) in the 2B region. During the growth of GSC progeny, fusomes become branched. The green bar indicates canonical Wnt signaling is detectable in the EC precursors of LL3 animals, and it becomes strongly activated in ECs at the mid-pupa stage, persisting into the adult fly. **(B-I)** L1 (B), L2 (C) and late-L3 gonads (D and E) with *fz3RFP* in B-D (gray, canonical Wnt signaling reporter), *3GRH4TH-GFP* in E (gray, canonical Wnt signaling reporter), Vasa (green in B-D, red in E, PGCs) and Tj (red in B-D, blue in E, ICs). B’-D’ and E only show *fz3RFP* and *3GRH4TH-GFP* channel, respectively. Dashed circles outline the gonad; dashed line marks a forming TF. **(F-I)** Early-pupa (F), Mid-pupa (G) and adult day 1 germaria (H and I) with *fz3RFP* in F-H (gray, canonical Wnt signaling reporter), *3GRH4TH-GFP*in I (gray, canonical Wnt signaling reporter), Vasa (green in F and G, germ cells), Tj (red in F and G, EC and follicle cell nuclei), LamC (red in H and I, TF and cap cell nuclear envelopes), and Hts (red in H and I, fusomes). Insets in F and G show enlarged images from the corresponding dashed squares in F and G. Arrowheads in F inset show ECs with *fz3RFP*. The genotype in F is *nos>gfp^RNAi^.* Asterisks in H and I mark GSCs. Scale bars are 10 μm.

In this study, we report that canonical Wnt signaling functions in the formation of ECs and maintains their long cellular protrusions to prevent Dpp and Wg signaling to GSC progeny, which would interfere with their proper differentiation. In germaria bearing ECs with blunted cellular protrusions, Dpp and Wg leak from the soma into the germ cell region. In the germ cells of these germaria, Dpp activates Dpp stemness signaling and Wg activates canonical Wnt signaling to promote transcription of CycB3, a G2/M regulator [29], and disrupts germ cell differentiation. Our results not only document a role for canonical Wnt signaling in EC formation during ovary development, but the findings also demonstrate a functional requirement for a somatic cell permeability barrier to prevent morphogen signals from reaching germ cells.

## Results

### The developing ovarian soma displays active canonical Wnt signaling

A previous study reported that canonical Wnt signaling is undetectable in the late larval ovary, but it is activated in germarial ECs of late pupae [16]. This finding was made using the *fz3RFP* reporter, which consists of the promoter region of *frizzled3* (*fz3*) followed by an open reading frame for red fluorescent protein (RFP) [30, 31]. Since a complete assessment of canonical Wnt signaling activation in the ovary throughout development was still lacking, we first examined the *fz3RFP* expression in the ovary at different developmental stages. In the L1 gonad (Fig. 1B and B’), *fz3RFP* was not detectable, but its expression was clearly observable in ICs of the L2 gonad (Fig. 1C and C’). In the late-L3 gonad (Fig. 1D and D’), *fz3RFP* was weakly expressed in developing TFs but more highly expressed in a subset of ICs and basal cells. Similar patterns were also observed in late-L3 gonads (Fig. 1E and E’) when using another canonical Wnt signaling reporter, *3GRH4TH-GFP,* which contains three Grainyhead (GRH) and four classic HMG-Helper (4TH) site pairs followed by green fluorescent protein (GFP) [32]. At the early-pupal stage, ICs in the germarium (considered to be ECs) expressed low levels of *fz3RFP* (Fig. 1F), while *fz3RFP* expression became strong in ECs at the mid-pupal stage (Fig. 1G) and remained in adult gemaria of newly eclosed flies (1-day-old) (Fig. 1H). Interestingly, in the 1-day-old adult germarium *3GRH4TH-GFP* was expressed only in one or two ECs that were in direct contact with GSCs (Fig. 1I), although a previous report showed *3GRH4TH-GFP* expression in posterior ECs and follicle stem cells [33]. Even so, our results indicate that canonical Wnt signaling is active in the developing ovarian soma and becomes highly active in germarial ECs from the mid-pupal stage.

### Canonical Wnt signaling in the developing soma controls EC formation and is required for proper germ cell differentiation

In the canonical Wnt signaling pathway, binding of Wnt ligands to Fz receptors leads to recruitment of Dsh and consequent disruption of the Axin destruction complex. This action stabilizes Arm, which subsequently enters the nucleus and interacts with Pygo (co-activator) and TCF to regulate expression of the downstream targets (Fig. 2A)[34]. It has been previously concluded that canonical Wnt signaling does not play a role during ovary development, as very low *fz3RFP* expression was detected in the larval ovary [16]. In addition, *arm* knockdown throughout development only causes a very mild increase in the number of undifferentiated cells carrying round-shape fusomes (spectrosome-containing cells; SCCs) [11]. However, our data showed that *fz3RFP* and *3GRH4TH-GFP* expression is detectable in ICs. Furthermore, no canonical Wnt signaling components other than Arm had been tested for functional effects in the developing ovary. We thus individually disrupted *dsh*, *arm* and *pygo* expression in the ovarian soma throughout development and examined 1-day-old germaria. For this purpose, we used *UAS-RNAi* lines driven by *tj-GAL4*, which is expressed in ICs of the larval ovary (Supplementary Fig. 1) [20]. At the anterior tip of the control 1-day-old germarium (Fig. 2B), two GSCs can be identified by their fusomes, i.e., the membrane-enriched organelle (yellow arrows) adjacent to niche cap cells (cap cell-GSC junction is indicated by a solid line). Additionally, 1 ± 0.7 spectrosome-containing CBs were designated as SCCs (n = 15 germaria) (indicated by the asterisk in Fig. 2B).

**Fig. 2.**
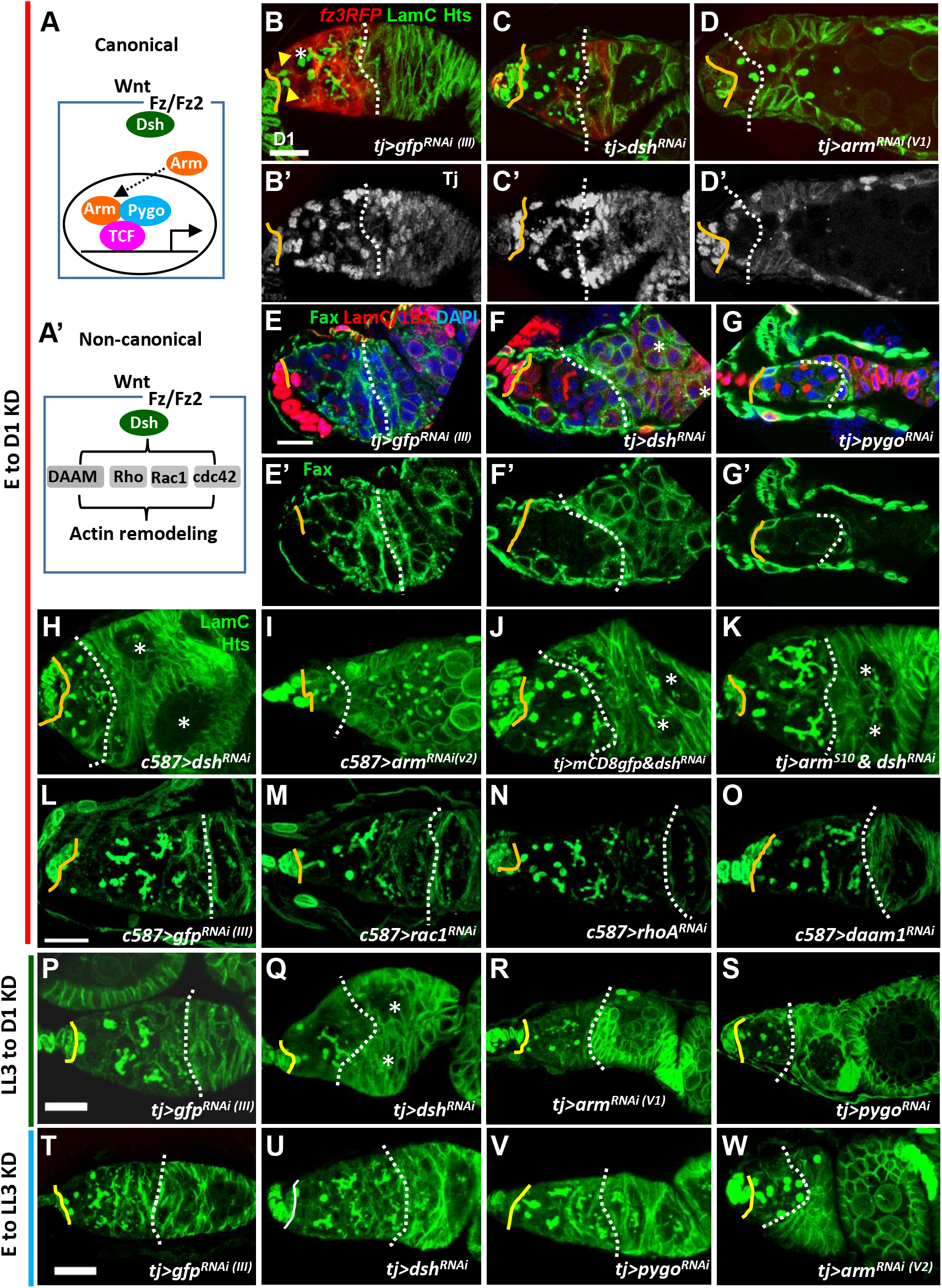
Canonical Wnt signaling in the developing soma controls escort cell formation and promotes germ cell differentiation. **(A and A’)** Canonical (A) and non-canonical Wnt signaling pathways (A’). Wnt signaling components were knocked down in the ovarian soma from embryo to adult day (D)1 (B-O), from late-L3 (LL3) to adult D1 (P-S), and from embryo to LL3 (T-W), and D1 germaria were examined. **(B-O)** *tj-GAL4>gfp^RNAi (III)^* (B and E)*, tj>dsh^RNAi^* (C and F), *tj >arm^RNAi (V1)^* (D), *tj>pygo^RNAi(V)^* (G), *c587>dsh^RNAi^* (H), *and c587>arm^RNAi(V)^* (I), *tj>mCD8-gfp & dsh^RNAi^* (J), *tj>arm^S10^ & dsh^RNAi^* (K), *c587>gfp^RNAi (III)^* (L), *c587>rac^RNAi^* (M), *c587>rhoA^RNAi^* (N) and *c587>daam1*^RNAi^ germaria (O) with LamC (green in B-D and H-O, red in E-G, terminal filament (TF) and cap cell nuclear envelopes), Hts (green in B-D and H-O, red in E-G, fusomes), Tj in B’ to D’ (gray, nuclei of cap, escort and follicle cells), Fax (green in E-G, EC membranes), and DAPI (blue in E-G, DNA). E’-G’ show only Fax channel. **(P-S)** *tj-GAL4>gfp^RNAi (III)^* (P), *tj>dsh^RNAi^* (Q), *tj >arm^RNAi (V1)^* (R) and *tj>pygo^RNAi^* germaria (S) with LamC (green) and Hts (green). **(T-W)** *tj-GAL4>gfp^RNAi (III)^* (T), *tj>dsh^RNAi^* (U), *tj>pygo^RNAi^* (N) and *tj >arm^RNAi (V2)^* germaria (W) with LamC (green) and Hts (green). Solid lines, GSC-cap cell junction; dashed lines, the 2A/2B boundary (or the junction between escort cells and follicle cells when 2A/2B boundary is missing). Arrowheads and an asterisk in Fig. 2B respectively mark GSCs and cystoblast. Two asterisks in F, H, K and Q denote two side-by-side egg chambers in the ovariole. Scale bars are 10 μm; B-D and H-K, E-G, L-O, P-S, and T-W have the same scale bar.

Differentiating germ cells containing branched fusomes were located posterior to the GSCs and wrapped by EC protrusions (marked by *fz3RFP*; Fig. 2B). Between cap cells (indicated by a yellow line) and the 2A/B boundary (indicated by a dashed line, Fig. 2B’), we observed 26.2 ± 3.3 ECs (n = 21 germaria) expressing Traffic jam (Tj, a Maf transcription factor [35]). In contrast, the *dsh*-and *arm*-knockdown (KD) germaria (Fig. 2C and D), fewer cysts with branched fusomes were found posterior to the GSCs, and this decrease in cysts was accompanied by SCC accumulation [*dsh*-KD, 4.7 ± 1.9 SCCs (n = 14 germaria), *P* < 0.001; *arm*-KD, 6.6 ± 4.6 SCCs (n = 14 germaria), *P* < 0.001; some SCCs were even observed within egg chambers]. As evidence for the disruption of Wnt signaling activity, *fz3RFP* expression in ECs was dramatically decreased in *dsh*- and *arm*-KD germaria (Fig. 2B-D). *dsh*- and *arm*-KD germaria also carried fewer ECs (*dsh* KD, 9.0 ± 3.8 ECs (n = 21); *arm* KD, 6.8 ± 4 ECs (n = 15), *P* < 0.001), resulting in shortened EC regions (space between cap cells and the 2A/B boundary, or space between cap cells and follicle cells if the 2A/B boundary is lost) (Fig. 2C’ and D’). In addition, Failed axon connections (Fax)-labeling was used to mark EC protrusions [36]. The germ cells in the control germarium were completely wrapped by ECs (Fig. 2E and E’), whereas the wrapping was disrupted in *dsh*-KD germaria (Fig. 2F and F’). Similar results could be obtained by disrupting *pygo* expression (Fig. 2G and G’) or by using another somatic GAL4 driver, *c587-GAL4* [37], to drive expression of *dsh^RNAi^* or another independent *arm^RNAi^* throughout developmental stages (Fig. 2H and I). Moreover, compared to *dsh*-KD germaria (Fig. 2J), overexpressing a constitutively active form of Arm, *arm^S10^* [38], in the *dsh*-KD soma throughout development expanded the EC region, decreased SCC accumulation [63% of *dsh*-KD & *mCD8gfp* germaria (n = 26) carrying more than 4 SCCs vs. 0% of *dsh*-KD & *arm^S10^* germaria (n = 20) carrying more than 4 SCCs, *P* < 0.001], and increased cyst cells bearing a branched fusome [0% of *dsh*-KD & *mCD8gfp* germaria (n = 26) carrying more 4-6 16-cell cysts vs. 95% of *dsh*-KD & *arm^S10^* germaria (n = 20) carrying 4-6 16-cell cysts, *P* < 0.001] (Fig. 2K). Given that these phenotypes were observed after knockdown of nearly every canonical Wnt signaling component, we concluded that canonical Wnt signaling is required for formation of ECs and their protrusions during ovary development, and that promotes germ cell differentiation

In the non-canonical Wnt signaling pathway, Wnt-Fz signals through Dsh and downstream effectors, such as Dsh Associated Activator of Morphogenesis 1 (DAAM1), Ras homolog gene family member A (RhoA) and Rac1, to modulate Actin remodeling, cell movement and division (see Fig. 2A’)[5]. Indeed, the occurrence of side-by-side-positioned cysts or egg chambers in *dsh*-KD germaria (21%, n = 19 germaria) was not rescued by expression of Arm^S10^ (25%, n = 20 germaria) (Fig. 2J and K, marked by asterisks); this phenotype was also present in *wnt*4-KD germaria (see Fig. 3D), supporting a role for Wnt4-mediated non-canonical Wnt signaling in the soma during ovary development [16]. Notably, disruption of downstream effectors of non-canonical Wnt signaling components in the ovarian soma throughout development did not cause shortened germaria (Fig. 2L-O). In agreement with a previous report [16], *rac1*-, *rhoA*-and *daam*-KD germaria displayed low levels of SCC accumulation (Fig. 2M-O); however, only a small fraction of germaria carried more than 4 SCCs [*rac1*-KD, 9.5% (n = 21); *rhoA*-KD, 33.3% (n = 21); and *daam*-KD, 15% (n = 20) of examined germaria; none were significantly different than controls, 14.3% (n = 21)]. These results suggested that canonical Wnt signaling is indispensable for specifying ECs, which promote germ cell differentiation.

**Fig. 3.**
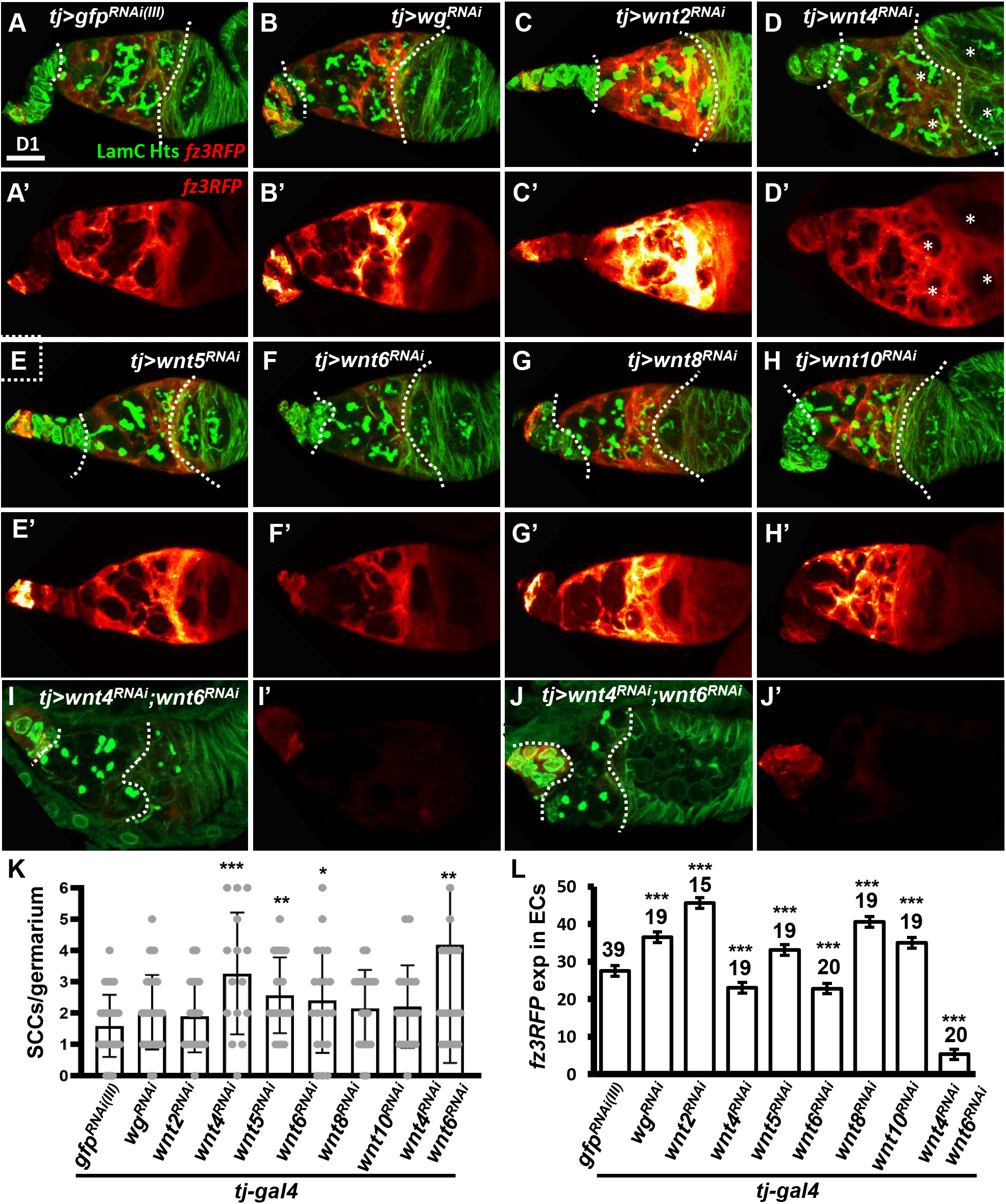
Wnt4 and Wnt6 activate canonical Wnt signaling in escort cells. **(A-H)** One-day (D)-old *tj>gfp^RNAi (III)^* (A), *tj>wg^RNAi^* (B), *tj>wnt2^RNAi^* (C), *tj>wnt4^RNAi^* (D), *tj>wnt5^RNAi^* (E), *tj>wnt6^RNAi^* (F), *tj>wnt8^RNAi^* (G), *tj>wnt10^RNAi^* germaria (H) and *tj>wnt4^RNAi^ &wnt6^RNAi^* (I and J) with *fz3RFP* (canonical Wnt signaling reporter), LamC (green, terminal filament and cap cell nuclear envelopes), and Hts (green, fusomes). A’-J’, Heatmap showing relative intensity, red = low to yellow = high, of *fz3RFP* in the germaria. Images were processed with Zen blue and Fire look-up table (LUT); color gradient bar indicates strength of *fz3RFP* from low (L, red) to High (L, gold). Scale bar, 10 μm; A-H have the same scale bar, K and L have the same scale bar. **(K)** Number of spectrosome-containing cells (SCCs) per germarium for indicated genotypes. **(L)** Expression of *fz3RFP* in the escort cell region (between cap cells and follicle cells; marked by dashed lines) of germaria with the indicated genotypes. *RNAi* was expressed throughout development until dissection. Error bar, mean ± SD. Statistical analysis, One-way ANOVA, *, *P* < 0.05; **, *P* < 0.01; ***, *P* < 0.001.

To understand the temporal requirement of somatic canonical Wnt signalling during ovary development, we knocked down *dsh*, *arm* and *pygo* at different developmental times by modulating GAL4 activity with GAL80^ts^; in these lines, GAL4 activity is suppressed at 18℃ but not at 29℃ [39]. Knockdown of *dsh*, *arm* and *pygo* from late-L3 to adult stage caused similar phenotypes, which were less severe than those seen with continuous knockdown (Fig. 2P-S). Suppressing *dsh* and *pygo* in the larval soma before the late-L3 stage did not result in aberrant germaria (Fig. 2T-V), while *arm*-KD germaria exhibited SCC accumulation (Fig. 2W), likely due to a requirement for the interaction between Arm and E-cadherin to promote PGC-IC intermingling (Supplementary Fig. 2) [20]. Taken together, these results show that canonical Wnt signaling acts on ICs, probably as early as late-L3, to maintain the EC population and promote germ cell differentiation.

### Wnt4 and Wnt6 in the developing soma stimulate canonical Wnt signaling in ECs

Five Wnts are known to be expressed in adult germarial somatic cells. Wg and Wnt6 are highly expressed in cap cells, while Wnt2 and Wnt4 are expressed in both cap cells and ECs [12, 13]. Wnt5 is also expressed at low levels, according to *RNA*-seq analysis of isolated ECs [12]. To determine which Wnt activates the somatic canonical Wnt signaling required for EC formation and germ cell differentiation, we individually knocked down Wnt ligands throughout development using *tj-GAL4* and examined germarial phenotypes and *fz3RFP* expression in 1-day-old ovaries. However, we did not observe overt phenotypes similar to those seen in *dsh*- or *arm*-KD (Fig. 3A-H). However, we did observe side-by-side cysts or egg chambers (indicated by asterisks in Fig. 3D and D’) after knockdown of *wnt4*, and we found slightly increased SCC numbers upon knockdown of *wnt4*, *wnt5* and *wnt6* (Fig. 3K). Somatic knockdown of *wg*, *wnt2*, *wnt5*, *wnt8* or *wnt10* increased *fz3RFP*, somatic knockdown of *wnt4* or *wnt6* decreased *fz3RFP* expression in ECs of 1-day-old germaria (Fig. 3L). Co-knockdown of *wnt4* and *wnt6* in the developing soma caused germaria to exhibit nearly absent *fz3RFP* expression, a shortened EC region, SCC accumulation (*wnt4 & 6* coKD germaria: 4.2 ± 3.7 SCCs, n = 22 germaria; *gfp*-KD germaria: 1.6 ± 1.0 SCCs, n = 20 germaria; *P* < 0.005), and side-by-side cysts or eggs (Fig. 3I-L), reminiscent of somatic *dsh*-KD germaria. We did not know why developmental knockdown of *wg*, *wnt2*, *wnt5*, *wnt8*, and *wnt10* would increase canonical Wnt signaling in ECs, but the effects illustrate the complexity of the Wnt signaling network. Nevertheless, our data suggested that Wnt4 and Wnt6 appear to be positive regulators of canonical Wnt signaling in ECs.

### Canonical Wnt signaling is activated in the germline when thickveins (Tkv) is suppressed in the soma during development

It has been reported that wrapping of germ cells by EC protrusions is crucial for proper germ cell differentiation; however, the reason for this requirement remains unclear. To specifically investigate the function of EC protrusions on germ cells, we knocked down Tkv (a Dpp receptor [24]) in the somatic cells during development. In this line, EC protrusions are disrupted, and SCC accumulation occurs, but the EC region is not shortened (Fig. 4A and B, and see Supplementary Fig. 7A and B)[37]. Intriguingly, although *RNA*-seq results showed an increased *fz3* mRNA transcript level in 1-day-old *c587>tkv^RNAi^* ovaries (Fig. 4C), *fz3RFP* expression was not increased in ECs of 1-day-old *tkv-*KD germaria, as compared to controls (see inset images in Fig. 4A and B). Co-knockdown of *dsh* and *tkv* in the developing ovarian soma did not prevent SCC accumulation but did cause a shortened EC region (Fig. 4D and E). These results indicate that canonical Wnt signaling in the soma is not involved in the impairment of germ cell differentiation in somatic *tkv*-KD germaria; instead, canonical Wnt signaling may be elevated within the germ cells of somatic *tkv*-KD germaria. We were not able to perform knockdown of Wnt signaling specifically in the germline of somatic *tkv*-KD germaria by the current genetic tools because the targeted signaling component would be knocked down both in the germline (by *nos-GAL4*) and in the soma (by *tj-GAL4*, which was used to knockdown *tkv*). Therefore, we knocked down Wnts in the developing soma of somatic *tkv*-KD ovaries. Somatic knockdown of *tkv* with *wnt2*, *wnt4*, *wnt5* or *wnt6* did not suppress SCC accumulation (Supplementary Fig. 3). Strikingly, simultaneous knockdown of *wg* and *tkv* in the developing soma partially rescued the germ cell differentiation defect, as evidenced by reduced SCC numbers, and increased 16-cell cyst numbers (Fig. 4F-H), indicating that Wg may activate canonical Wnt signaling in the germline of somatic *tkv*-KD germaria.

**Fig. 4.**
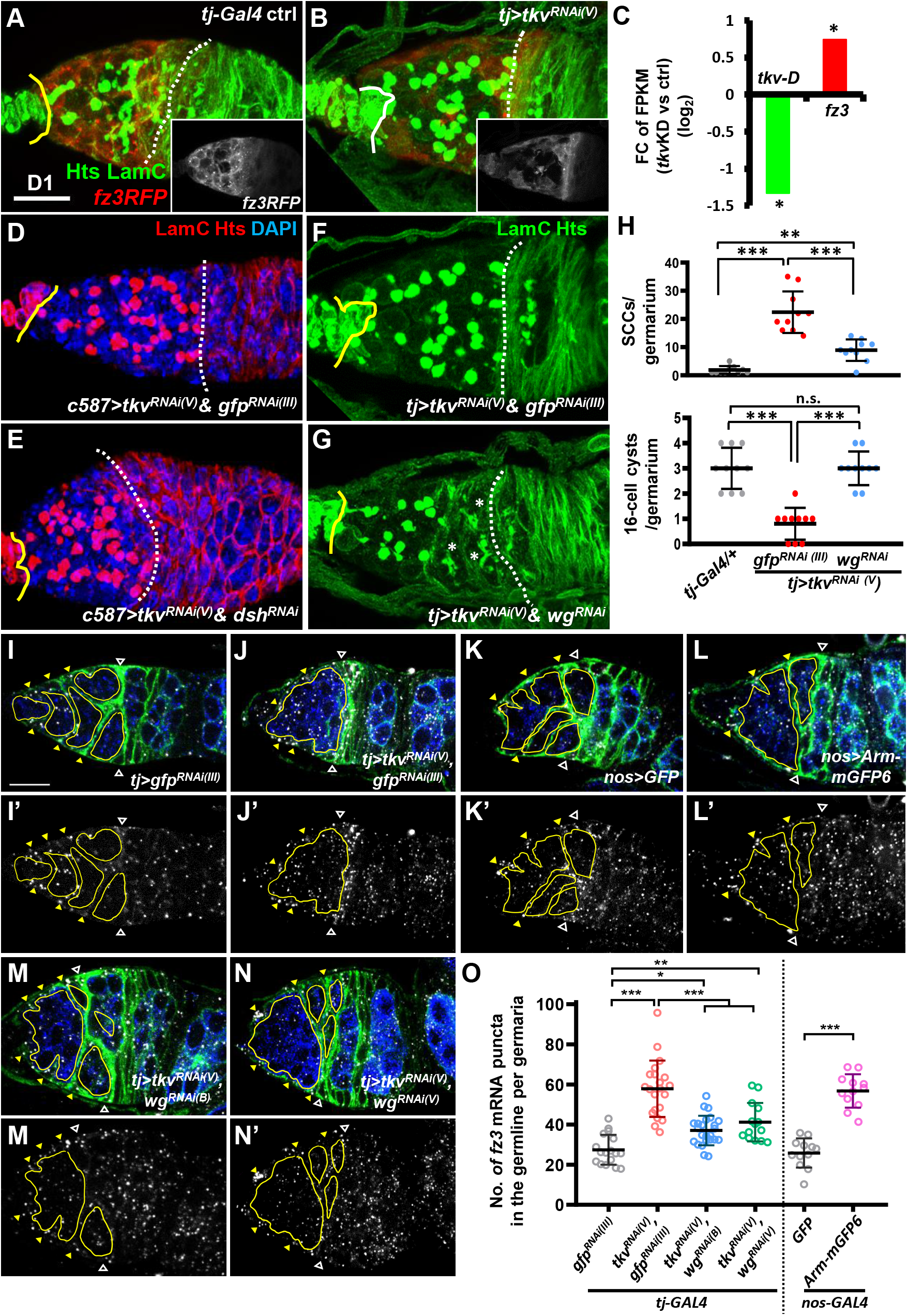
Wg signaling is activated in the germ cells of somatic-*tkv* knockdown germaria. **(A and B)** *tj-GAL4/+* (A) and *tj>tkv^RNAi (V)^* germaria (B) with LamC (green, TF and cap cell nuclear envelopes), Hts (green, fusomes) and *fz3RLFP* (red, canonical Wnt signaling reporter). Insets in A and B show only *fz3RFP* channel. **(C)** Fold-change (FC) of RNA-seq based gene expression values (log2) for *tkv* transcript variant D (*tkv-D*) and *fz3* in one-day (D)-old control (ctrl, *UAS-tkv^RNAi (N)^/+*) and *c587>tkv^RNAi (N)^* anterior ovarioles compared with *UAS-tkv^RNAi (N)^/+* (control, ctrl). FPKM, fragments per kilobase of transcript per million mapped reads. *, *P* < 0.05. Statistical analysis was performed with two biological replicates. **(D** to **G)** *c587>tkv^RNAi (V)^ & gfp^RNAi (III)^* (D), *c587>tkv^RNAi (V)^ & dsh^RNAi^* (E), *tj>tkv^RNAi (V)^ & gfp^RNAi (III)^* (F) and *c587>tkv^RNAi (V)^ & wg^RNAi^* germaria (G) with LamC (red in D and E, green in F and G), Hts (red in D and E, green in F and G) and DAPI (blue, DNA, in D and E). Yellow lines denote the junction between cap cell and GSC; dashed line mark the 2A/2B boundary or the junction between escort cells and follicles when the 2A/B boundary is missing. Asterisks mark 16-cell cysts. **(H)** Numbers of spectrosome-containing cells (SCCs) and 16-cell cysts per germarium of flies with the indicated genotypes. **(I-N)** *In situ* hybridized *tj>gfp^RNAi (III)^* (I), *tj>tkv^RNAi (V)^ & gfp^RNAi (III)^* (J), *tj>gfp* (K), *tj>arm-mGFP6* (L), *tj>tkv^RNAi (V)^ & wg^RNAi(V)^* (M) and *tj>tkv^RNAi (V)^ & wg^RNAi(B)^*germaria (N) with labeling for Fax (green, escort cell membrane extension), Vasa-GFP (blue, germ cells), and *fz3* mRNA (gray). I’-N’ show the *fz3* channel. Hollow triangles point to the 2A/B boundary; yellow triangles indicate escort cell region; germ cell regions before the 2A/B boundary are outlined by yellow circles. **(O)** Number (No.) of *fz3* mRNA puncta in the germline per germarium with the indicated genotypes. *, *P* < 0.05, **, *P* < 0.01; ***, *P* < 0.001. Error bars indicate mean ± S.D., One-Way ANOVA was used for statistical analysis. *RNAi* was expressed throughout development. Scale bar, 10 μm. A and B, D-G, and I to N are 3D-reconstructed images.

We did not detect *fz3RFP* expression in germ cells, perhaps because *fz3RFP* only effectively reports Wnt signaling in somatic cells. We thus examined *fz3* mRNA expression in somatic *tkv*-KD germaria by *in situ* hybridization. In the control germarium (Fig. 4I), *fz3* transcripts were detected in the cytoplasm of anterior germ cells and ECs, while *fz3* transcript levels were dramatically increased in the germ cells of somatic *tkv*-KD germaria (Fig. 4J and O). Of note, no *fz3* mutants or *UASp-RNAi* lines were available to test the specificities of *fz3* anti-sense probes we used. Instead, we overexpressed *arm* or knocked down *axin* in the germline to force canonical Wnt signaling activation, and then we examined *fz3* expression. We found that *fz3* transcripts were significantly increased in the germline with *arm* overexpression (Fig. 4K, L and O), and in *axin*-KD germ cells (Supplementary Fig. 4). As expected, increased *fz3* transcripts in the germline of somatic *tkv*-KD germaria could be suppressed by knockdown of *wg*, using two independent *RNAi* lines (Fig. 4M-O). These results suggested that Wg from the soma contributes to germ cell differentiation defects in somatic *tkv*-KD germaria. It is likely that when *tkv* is disrupted in the soma, the receipt of Wg by germ cells activates canonical Wnt signaling, which is deleterious for germ cell differentiation.

### Decreasing CycB3 expression in the germline of somatic tkv-KD germaria partially rescues germ cell differentiation

From the *RNA*-seq data, we noticed that transcripts of some Cyclins (e.g., CycB, CycB3, and CycE) and Cyclin-dependent protein serine/threonine kinase regulators (e.g., CycT) were significantly increased in 1-day-old somatic *tkv-*KD germaria (Fig. 5A). Among the increased transcripts, CycB, CycB3 and CycE are known to be required for GSC maintenance [40–42], and increased CycB3 expression delays CB differentiation [42]. Given the continuous proliferation of SCCs in somatic *tkv*-KD germaria (Supplementary Fig. 5), and since an antagonism between cell cycle regulators and differentiation genes has been proposed [43], we next asked if reducing Cyclins could suppress germline differentiation defects in somatic *tkv*-KD germaria. We individually knocked down *cycB*, *cycB3* and *cycE* in the germline using *nos-GAL4*, along with *tkv* knockdown in the soma using *tj-GAL4* throughout developmental stages (Fig. 5B-E). Of note, the *tkv^RNAi(V)^* line used in this study was not effectively expressed in the germline due to its *UASt* promoter (see Fig. 4H and 5F), and therefore the role of Tkv in the germline is uncertain with regard to GSC maintenance. Knockdown of *cyclins* in the developing soma did not cause obvious defects in germaria, but knockdown of *cycB* and *cycE* in the germline throughout development respectively caused GSC loss and SCC accumulation (Supplementary Fig. 6). Furthermore, knockdown of *cycB* in the germline of somatic *tkv*-KD germaria did not rescue the germ cell differentiation defect (Fig. 5B-F and F’), and germ cells were completely lost from *tj&nos>tkv^RNAi^&cycE^RNAi^* germaria, which displayed a thin tubular-like shape (Fig. 5D). Strikingly, knockdown of *cycB3* in the germline of somatic *tkv*-KD ovaries decreased SCCs and increased 16-cell cysts (Fig. 5E, F and F’), and it partially rescued EC protrusions (Supplementary Fig. 7). These effects suggested that the increase of CycB3 in the germline suppressed germ cell differentiation in the somatic *tkv*-KD germaria.

**Fig. 5.**
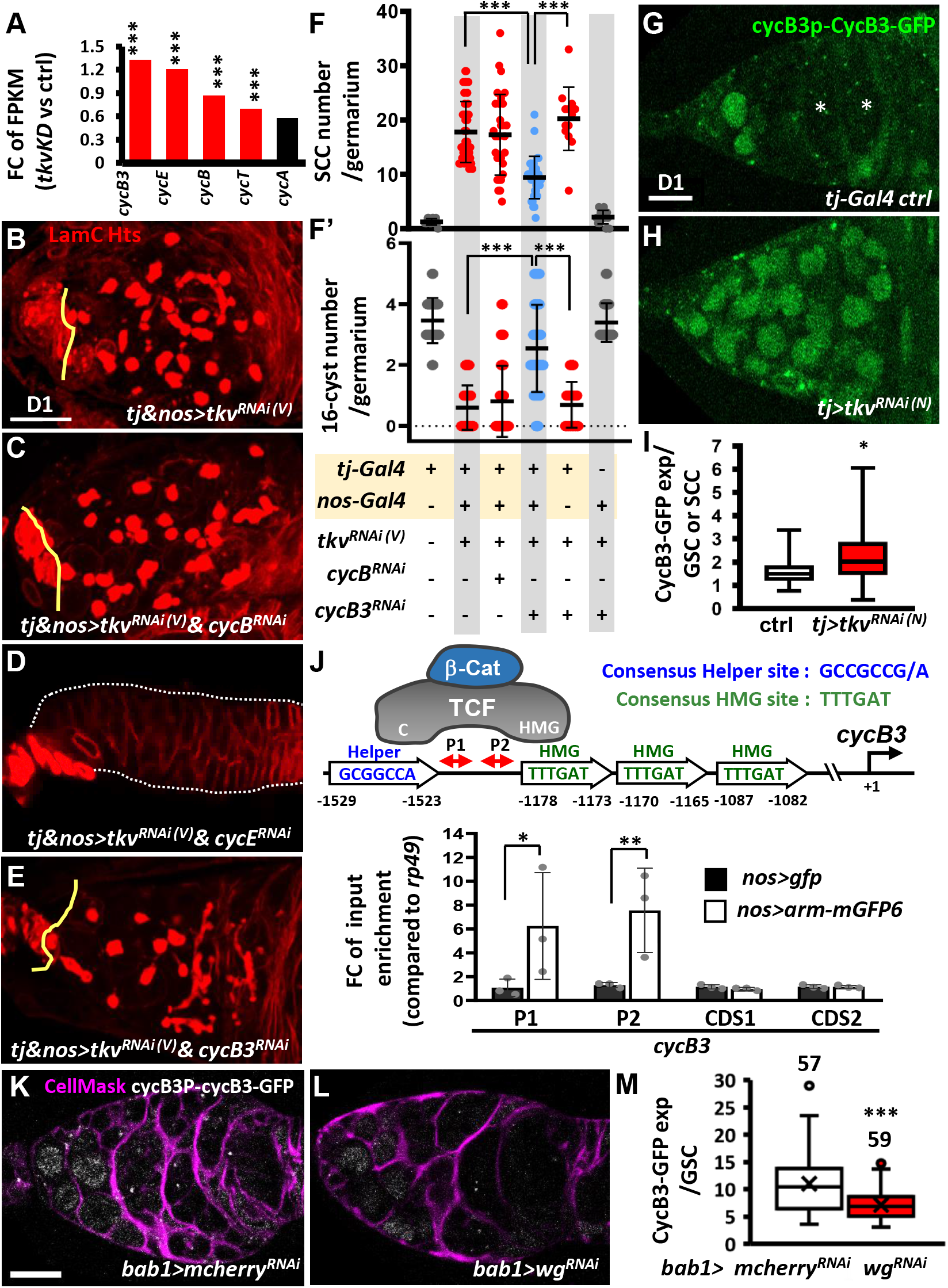
Canonical Wnt signaling transcriptionally activates *cycB3* to suppress differentiation in the germline of somatic-*tkv* knockdown germaria. **(A)** Fold-change (FC) of *RNA*-seq-based gene expression values (log2) for indicated *cyclin (cyc)* gene in one-day (D)-old control (ctrl, *UAS-tkv^RNAi(N)^/+*) and *c587>tkv^RNAi(N)^* anterior ovarioles compared with *UAS-tkv^RNAi(N)^/+* (control, ctrl). FPKM, fragments per kilobase of transcript per million mapped reads. ***, *P* < 0.001. Statistical analysis was performed with two biological replicates. **(B to E)** One-day-old *tj & nos>tkv^RNAi (V)^*(B), *tj & nos>tkv^RNAi(V)^ & cycB^RNAi^* (C), *tj & nos>tkv^RNAi(V)^ & cycE^RNAi^* (D) and *tj & nos>tkv^RNAi (V)^ & cycB3^RNAi^* (E) and with LamC (red, terminal filament and cap cell nuclear envelopes) and Hts (red, fusomes). Solid lines mark junction between GSCs and cap cells; dashed lines outline the germaria in D. **(F and F’)** Numbers of spectrosome-containing cells (SCCs) (F), and 16-cell cysts per germarium (F’) of flies with the indicated genotypes. **(G and H)** Live image of 1-day-old *tj>GFP^RNA (III)^* (G) and *tj>tkv^RNAi (V)^* germaria (H) bearing *cycB3P-cycB3-gfp* (Green, CycB3-GFP). **(I)** Box plot shows expression of CycB3-GFP in GSCs or SCCs in the indicated genotypes. **(J)** Schematic shows how β-catenin (β-Cat) interacts with TCF, which binds to HMG and Helper sites of the *cycB3* promoter through its HMG and C domains, respectively. ChIP analysis of TCF binding in 1-day-old ovaries; the chromatin from *nos>gfp* and *nos>arm-mgfp6* cells was precipitated with GFP-Trap beads. Co-precipitated DNA was analyzed by qPCR using two sets of primers (P1 and P2) against the region between Helper and HMG sites. The amplicons of two different coding regions were used as negative controls. **(K and L)** One-day-old *bab1>mcherry^RNAi^* (K) and *bab1*>*wg^RNAi (V)^*(L) with cycB3P-CycB3-GFP (gray) and CellMask (Magenta, cell membrane). (M) Average of CycB3-GFP expression in GSCs of indicated genotypes. Number of GSCs analyzed is shown above each bar. Differences in F and F’ were analyzed by one-way ANOVA; data in I and M were analyzed by Student’s *t* test, and in J were analyzed by two-way ANOVA. Solid line in the box of I and M is median; cross in M is Mean. Error bars represent SD; *, *P* < 0.05; **, *P* < 0.01; ***, *P* < 0.001. *RNAi* was expressed throughout development until dissection. Scale bar is 10 μm.

### Wg signaling promotes CycB3 expression in SCCs upon somatic knockdown of tkv

We next examined CycB3 expression in live germaria, using a *cycB3* promoter *(P)*-CycB3-GFP transgene [42]. We used this approach due to a lack of anti-CycB3 antibody and high background from anti-GFP staining. In the control germarium (Fig. 5G and I), CycB3-GFP was expressed in GSCs, germ cells posterior to GSCs, and some follicle cells, but it was absent in late-differentiating cysts (marked by asterisks). In somatic *tkv*-KD germaria, CycB3-GFP expression was further enhanced in GSCs and prospective SCCs (Fig. 5H and I), which were identified by the nucleus size being comparable to control GSCs. These results raise the possibility that canonical Wnt signaling may promote CycB3 expression at the transcriptional level.

Activation of Wnt signaling induces β-catenin nuclear translocation and interaction with TCF, turning on target gene transcription [2]. We found that the *cycB3* promoter includes a putative Helper-HMG pair element (Helper: -1529-1529bp; HMG: -1178-1173bp) (Fig. 5J); Helper and HMG sequences are respectively recognized by the C-clamp and HMG domain of TCF [44]. We used chromatin immunoprecipitation (ChIP) to examine whether the Arm (tagged with GFP)-TCF complex binds to the Helper-HMG pair element of the *cycB3* promoter in 1-day-old ovaries carrying *nos>arm-gfp*. To determine the Arm-TCF complex occupancy on the Helper-HMG pair element, we used qPCR to amplify two fragments (P1 and P2) located in the promoter region between the Helper and HMG sites (Fig. 5J). The amounts of amplified P1 and P2 fragments from *nos>arm-gfp* ovaries were 6-to 7-fold higher than those from *nos>gfp* ovaries, while the amounts of PCR product amplified from the coding region of *cycB3* in *nos>gfp* ovaries showed no difference (Fig. 5J). Furthermore, knockdown of *wg* using another somatic driver (*bab1-GAL4*, which is expressed in ICs and enriched in cap cells where Wg is generated after pupal stages [37], significantly reduced *cycB3-GFP* expression (Fig. 5K-M). These results indicate that *cycB3* is a novel target of canonical Wnt signaling. Taken together, the data suggests that when *tkv* expression is disrupted in the developing ovarian soma, germline canonical Wnt signaling is upregulated and transcriptionally promotes expression of CycB3, which in turns suppresses germ cell differentiation.

### Wnt-cycB3 regulation in the germline promotes germ cell differentiation in normal germaria

To determine if enhanced Wnt signaling promotes SCC accumulation via loss of somatic Tkv function in the soma, we directly overexpressed Arm or suppressed Axin (a negative regulator of Wnt signaling [45]) in the germline throughout development and examined 1-day-old germarial phenotypes. These germaria did not exhibit SCC accumulation, but we observed increased *cycB3-GFP* expression and higher numbers of 16-cell cysts located in the anterior germarium compared with the numbers of 16-cell cysts present in region 2 of the control germaria (Supplementary Fig. 8A-I); these findings were in agreement with a previous study [46]. In addition, disruption of canonical Wnt signaling in the germline was previously shown to slightly increase SCC numbers, suggesting a delay of CB differentiation [46]. Furthermore, knockdown of *cycB3* in the germline of *nos>axin^RNAi^* germaria decreased 16-cell cysts (Supplementary Fig. 8J-L). These results further confirm the existence of a Wnt signaling-CycB3 regulatory axis that is important for germline homeostasis. However, somatic cells appear to play a direct or indirect role in promoting or suppressing germ cell differentiation by Wnt signaling-CycB3 regulation, at least in part through the action of Tkv.

### Blunted EC protrusions allow Wg and Dpp to signal in the germline

We next asked how Wg-mediated canonical signaling becomes activated in the germline of somatic *tkv*-KD germaria. Previous reports showed that Wg is produced from cap cells [13, 47–49] and received by follicle stem cells to promote their maintenance and proliferation [48, 49]. EC protrusions wrap germ cells [50] and are disrupted in somatic *tkv*-KD germaria (see Supplementary Fig. 7)[37], raising the possibility that Wg might be normally restricted to somatic cells due to EC protrusions. Therefore, Wg may be able to access germ cells that are not wrapped by EC protrusions. To test this hypothesis, we used GFP-Wg in which GFP is inserted into the *wg* locus [51] to examine the distribution of Wg in live 1-day-old control and somatic *tkv*-KD germaria labeled with CellMask, a cell membrane dye. GFP-Wg granule numbers in TF and cap cells (19.6 ± 3, n = 12), where Wg is produced, were decreased when *wg* (9.1 ± 3, n = 17, *P* < 0.001) or *gfp* (7.8 ± 4.3, n = 14, *P* < 0.001) were knocked down in the developing soma using *c587-GAL4* (Fig. 6A and B, and Supplementary Fig. 9), demonstrating that GFP-Wg expression can fairly represents Wg expression. GFP-Wg expression was not altered in somatic *tkv*-KD ovaries as compared to control (Fig. 6C), while GFP-Wg distribution was altered (Fig. 6D-F). In the control germarium (Fig. 6E-E’’ and Supplementary Fig. 10), GFP-Wg was mainly present in cap cells (arrows), and it was observed in ECs (indicated by yellow asterisks) as well as in follicle cells and germ cells posterior to the 2A/2B boundary; very few GFP-Wg signals were observed in the germ cell zone before the 2A/2B boundary (2.3 ± 1.2 granules, n=11 germaria). In contrast, in addition to above-mentioned somatic cells and germ cells after follicle cell layer (2A/B boundary was missing), GFP-Wg signals were increased in the germ cell zone before the follicle cell layer of the somatic *tkv*-KD germarium (10.9 ± 2.3 granules, n= 12 germaria; *P*<0.001) (Fig. 6F-F’’ and Supplementary Fig. 10). We further confirmed this explanation by examining the Wg distribution in the *bag-of-marbles* (*bam*) mutant germarium, which display blunted EC protrusions (Supplementary Fig. 11B) due to defective germ cell differentiation [28]. The GFP-Wg distribution in the germline zone before follicle cell layer (2A/B boundary was missing) of the *bam* mutant was similar to the distribution in the somatic *tkv*-KD germaria (Fig. 6G-G’’ and Supplementary Fig. 10), although GFP-Wg expression seemed to be increased. Consistent with this observation, CycB3-GFP expression was also increased in SCCs of *bam* mutant germaria (Supplementary Fig. 11). ECs did not produce *wg* transcripts (Supplementary Fig. 12), indicating that niche-produced GFP-Wg distributed in the germline of somatic *tkv*-KD germaria is due to the lack of EC protrusions.

**Fig. 6.**
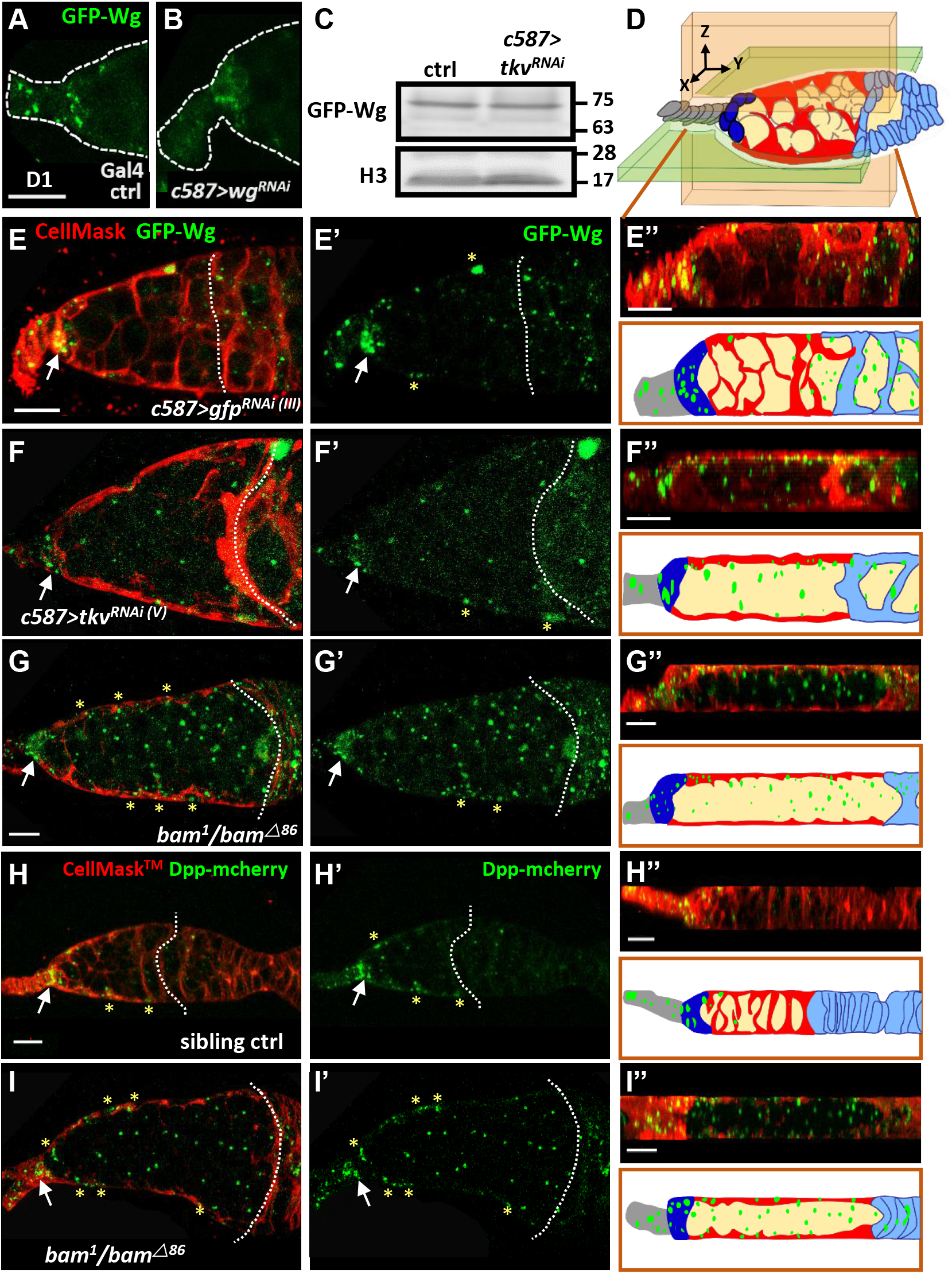
Cellular protrusions of escort cells serve as a physical permeability barrier to prevent Wg and Dpp distribution in the germline. **(A and B)** Live images of anterior part of one-day (D)-old GAL4 control (ctrl) (A) and *c587>wg^RNAi^* germaria (B) bearing GFP-wg (green). Dashed lines show the edge of the germarium. **(C)** Representative immunoblot shows that Wg-GFP expression (anti-GFP antibody) is similar in one-day (D)-old control (ctrl) and *c587>tkv^RNAi (V)^* ovaries. Histone (H3) was used as a loading control. Molecular weight markers are indicated to the right of the blots. **(D)** Schematic of a germarium with a three-axis (X, Y, and Z) coordinate system. The directions of each axis are shown. The Y-axis is defined as anterior to posterior. An XY section is shown as the light green shaded area, and an XZ section is shown as a light pink shaded area. Terminal filament cells, gray; cap cells, dark green; escort cells, red; germ cell, light yellow; follicle cells, light blue. **(E-I)** Live images of 1-day-old control (E), *c587>tkv^RNAi(v)^* (F), and *bam^1^/bam^△86^* mutant (G and I), sibling control germaria (H), bearing *GFP-wg* (green in E-G), Dpp-mcherry (green in H and I) and labelled with CellMask (red, cell membrane). E’’-I’’ are optical sections in the XZ plane; the corresponding schematic with the cell types is shown in D and corresponds to E’’-I’’; green color shows the distribution of Wg (E”-G”) or Dpp (H’’-I’). Scale bar, 10 μm; A and B, E and F, and H and I share the same scale bar. Germaria were examined for wg-GFP distribution in control (n = 12), *c587>tkv^RNAi(v)^* (n = 25), and *bam^1^/bam^△86^* mutant (n = 15). Germaria were examined for Dpp-mechrry distribution in the control (n = 8), and *bam^1^/bam^△ 86^* mutant (n = 12). Control genotypes in A, C and E are *c587>gfp^RNAi(III)^, GFP-wg/+*; in H is (*+/Dpp-mcherry; bam^1^or bam^△86^/+*). Arrows point to the cap cell region. Dashed lines in E and H mark the 2A/2B boundary, and F, G and H mark in the junction between escort cells and follicle cells (the 2A/2B boundary is lost). Asterisks mark GFP signals present in ECs.

We next wanted to know if EC protrusions also limit the distribution of Dpp (mammalian BMP; stemness factor that maintains GSC fate), which is produced from cap cells. To answer this question, we used mCherry-tagged Dpp (Dpp-mCherry), which was expressed in cap cells and ECs in the control germarium (Fig. 6H-H’’). Remarkably, Dpp-mCherry signal was spread throughout the germ cell zone of the *bam* mutant germarium (*bam* mutant; 28.9 ± 7 granules, n= 10 germaria vs. sibling control: 2.7 ± 2 granules, n= 18 germaria; *P*<0.001) (Fig. 6I-I’’ and Supplementary Fig. 13). These results suggest that EC protrusions wrap germ cells and prevent them from receiving cap cell-produced signals.

### EC protrusions act as a physical barrier to compartmentalize germ cells

We next directly tested if the germline is isolated from the external environment by EC protrusions using a previously developed permeability assay [52]. In this assay, ovaries were dissected and incubated in medium containing a fluorescently labeled 10-kDa dextran dye. The dye accessibility to germ cells was assessed. In the control germarium (n = 10 germaria)(Fig. 7A-A’), the fluorescence signal (black) overlapped with EC protrusions (red, marked by CellMask) but was excluded from germ cells. In contrast, in somatic *tkv*-KD germarium with blunted EC protrusions (n = 12 germaria)(Fig. 7B-B’’), the fluorescent dye was observed between all germ cells. This result suggests that EC protrusions isolate germ cells from the external environment.

**Fig. 7.**
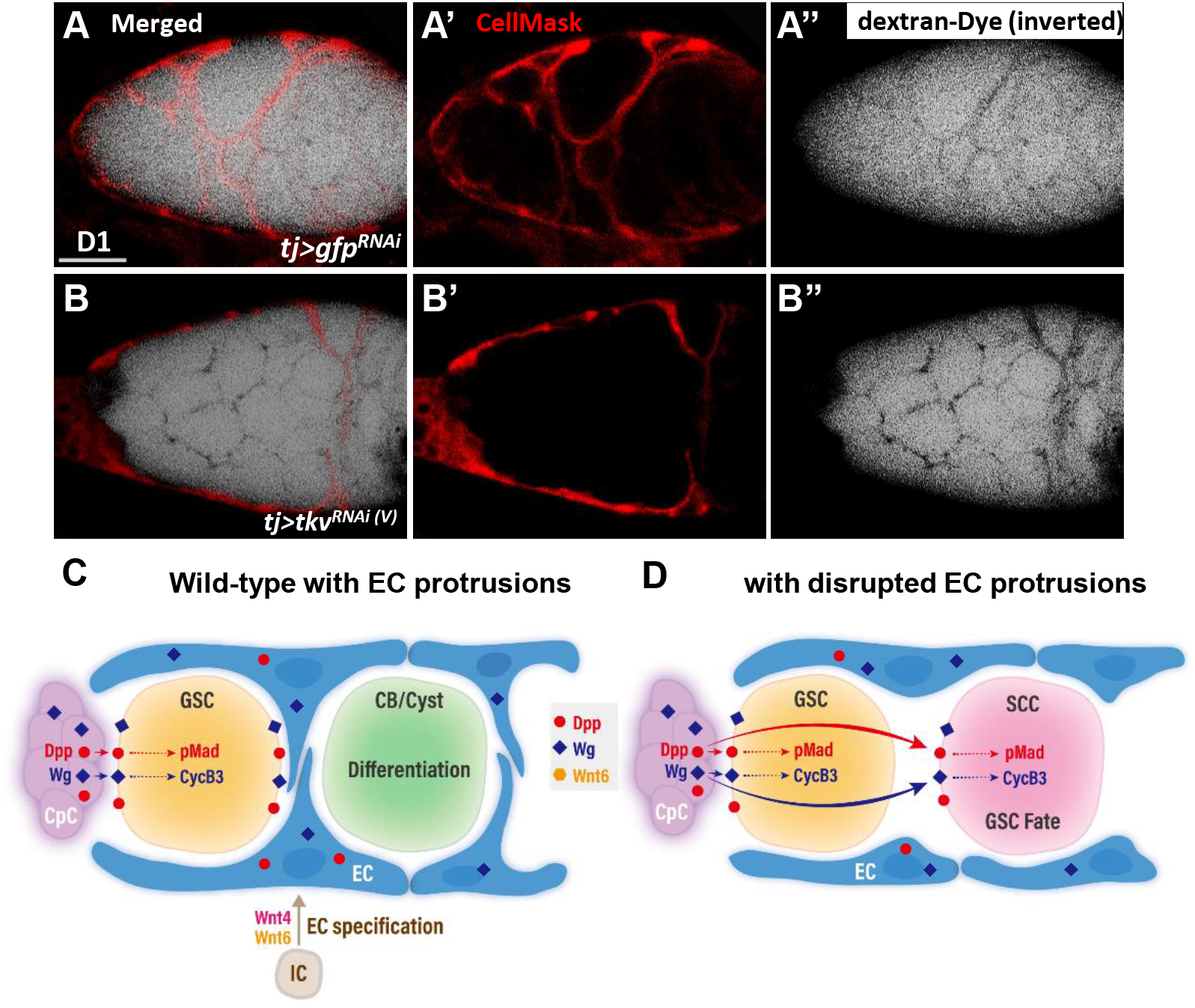
A permeability assay for the ovary assay reveals the role of EC protrusions warpping germ cells and acting as a physical permeability barrier. **(A and B)** One-day-old live *tj>gfp^RNAi^* (A) and *tj>tkv^RNAi(V)^* germaria (B) with CellMask (red, EC cell membrane) and dextran-488 (dextra signals are inverted for better visualization, a fluorescence dye). A and B are merged images; A’ and B’ show only CellMask channer and A’’ and B’’ show only dextran-488 channel. Scale bar, 10 μm. **(C and D)** Model of canonical Wnt signaling in EC specification and promotion of EC protrusions to set Dpp and Wg territories and maintain germline homeostasis. **(C)** In the developing wildtype ovary, canonical Wnt signaling in intermingled cells (ICs) is at least in part activated by Wnt4 and Wnt6. This signaling is critical for escort cell (EC) formation and maintains EC protrusions. The EC protrusions serve to compartmentalize GSC progeny and shield the germ cells from Dpp and Wg produced by cap cells (CpC), allowing the GSC progeny to properly differentiate. In GSCs, Dpp signaling leads to Mad phosphorylation (pMad), and upregulation of CycB3 probably occurs via transcriptional activation by Wg signaling. These events are critical to maintain GSC fate. **(D)** In the germarium with blunted EC protrusions, Dpp and Wg also signal to GSC progeny and disrupt their differentiation. CB, cytoblast; cyst, germ cell cysts.

## Discussion

Wnts are critical and conserved morphogens that control development of various organs. However, the nuanced roles of Wnt signaling in ovary development are still undefined, and how Wnt signaling territory is set within the tissue remains largely unclear. Here, we report that canonical Wnt signaling is activated in the ovarian ICs to promote EC formation and maintain EC protrusions. When canonical Wnt signaling is disrupted in ICs, EC number is decreased and EC cellular protrusions are disrupted. In the adult wild-type germarium (Fig. 7C), cap cells express Dpp and Wg signals, which respectively lead to Mad phosphorylation and transcriptional activation of CyCB3 in GSCs to maintain GSC fate. In addition, Dpp and Wg can signal to ECs as well (see Fig. 6) [13]. When EC protrusions are present, GSC progeny are compartmentalized, preventing aberrant activation of Dpp/Wnt signaling in GSC progeny to allow their proper differentiation. By contrast, without EC protrusions (Fig. 7D), Wg and Dpp are no longer restrained in the soma and diffuse to GSC progeny, where they suppress germ cell differentiation. Taken together, our data suggest that canonical Wnt signaling in the developing ovarian soma promotes the development of ECs, and the EC protrusions act as a physical permeability barrier to establish the territory of morphogens produced by the GSC niche. This barrier is necessary for proper differentiation of GSC progeny. Similar physical cell barriers to prevent the reception of morphogen signals by germ cells might exist in other organs and organisms, where they may act as determinants of tissue patterning.

### Both canonical and non-canonical Wnt signaling in the soma of larval gonads are required for germ cell differentiation

A switch between Wnt4-Dsh-mediated non-canonical and canonical Wnt signalling in larval gonads and adult germaria has been proposed (Upadhyay et al., 2018). In their study, Upadhyay and colleagues reported that two non-canonical Wnt signaling reporters were expressed in ICs of late larval gonads, but the reporter expression levels were decreased in adult ECs. Meanwhile, expression of the canonical Wnt signaling reporter, *fz3RFP*, was observed in a reverse pattern; it was not expressed in ICs, but it became strongly expressed in ECs. Knockdown of non-canonical Wnt signaling components decreased IC number and disrupted IC-PGC intermingling, resulting in a milder germ cell differentiation defects in adult germaria. In contrast, knockdown of non-canonical Wnt signaling components in adult ECs did not cause obvious defects.

In our study, we did not observe obvious defects in newly eclosed flies when canonical Wnt signaling was knocked down in the ovarian soma from embryonic to late-L3 stages, suggesting a dispensable role of canonical Wnt signaling in the soma before the late-L3 stage. However, we could detect the expression of two different canonical Wnt signaling reporters in ICs of L2 and late-L3 gonads. In addition, although Dsh is involved in both canonical and non-canonical Wnt signaling, we did not find IC-PGC intermingling defects when *dsh* was knocked down in the soma (see Supplementary Fig. 2C). Nevertheless, overexpressing a constitutively active form of Arm could not rescue the side-by-side cyst or egg chamber phenotype in somatic *dsh*-KD germaria (see Fig. 2F); this phenotype is also observed in *tj>wnt4^RNAi^* germaria (see Fig. 3D). Thus, both canonical and non-canonical Wnt signaling function in the larval soma, and the putative switch between non-canonical and canonical Wnt signaling in the ovarian soma might occur as early as the late-L3 stage.

### EC protrusions compartmentalize germ cells to block cap cell-produced maintenance cues

The boundaries of Wg signaling contribute to the patterning of various cell types in tissues throughout the organism. Several determinants of Wg territories have been reported. For example, glypicans (cell surface heparan sulfate proteoglycans) were shown to affect cell surface localization of morphogens [53], including Wg, Hedgehog (Hh) and Dpp. The fly has two glypicans, Dally and Dally-like protein (Dlp) [54]. Dally is expressed in cap cells to facilitate short-range Dpp trans signaling in GSCs [55, 56], while Dlp is expressed in ECs for Wg long-range travel from cap cells to follicle stem cells [48]. It has been proposed that Dally acts a classic co-receptor, while Dlp is like a gatekeeper, helping to transfer Wg from the source cells to distal cells [57]. Interestingly, overexpression of Dlp in ECs attenuates Wnt signaling and results in the absence of *fz3RFP* expression, fewer ECs, blunted EC protrusions, and defective germ cell differentiation [10]. On the other hand, knockdown of Dlp in ECs phenocopies Wnt overexpression, resulting in increased EC number and GSC loss without affecting germ cell differentiation [10]. These results suggest that Dlp and Fzs trap Wg, at least partially on the EC surface. However, this explanation cannot fully account for the inactivation of canonical Wnt signaling in germ cells, since germ cells are closely associated with ECs and express low levels of Fzs, according to single cell-sequencing results from the larval gonad [58] and adult ovaries [59]. In addition, somatic *tkv*-KD germaria do not show decreased *fz3RFP* expression, obvious changes in EC number, or GSC loss [37], suggesting that Wg-trapping molecules are still expressed on the EC surface. In this study, we showed that GFP-Wg is distributed in the germ cell zone of somatic *tkv*-KD or *bam* mutant germaria, indicating that the expansion of GFP-Wg territory does not rely on ECs themselves. Therefore, the EC protrusions may generate a compartment to keep germ cells shielded from Wg. In such case, the interfaces between ECs and germ cells would not encounter Wg, while the outer surfaces of ECs (facing sheath cells) would have the opportunity to trap cap cell-secreted Wg. A similar mechanism seems to restrict Dpp in GSCs. In addition to Wg, cap cells also produce Hh, Wnt2, Wnt4 and Wnt6 [13, 20]. Unfortunately, we do not have tools available to investigate whether the distributions of these molecules are altered when EC protrusions are blunted. In addition, we do not know how germ cells remain unresponsive to Wnt2 and Wnt4, which are produced by ECs [12, 13]. Perhaps ECs display a cell polarity that causes Wnt2 and Wnt4 to be secreted only from the outer surface. Despite these remaining uncertainties, our results show that depletion of Wg alone can partially rescue germ cell differentiation defects in somatic *tkv*-KD germaria. Furthermore, the results suggest thatEC protrusions physically wrap GSC progeny to block receipt of cap cell-derived maintenance cues, allowing germ cells to undergo proper differentiation, in line with the hypothesis made by Banisch and colleagues [60].

### Cell barriers in setting morphogen territories may be evolutionary conserved

Soma-germline interactions are critical for germ cell differentiation, and in the fly ovary, EC-germline interactions are particularly important for germ cell differentiation. The long cellular protrusions of ECs wrap germ cells [28, 60], and disruption of these protrusions causes germ cell differentiation defects. Conversely, blocking germ cell differentiation also impairs EC protrusions (see Fig. 6G and I) [28]. In this study, we show that protrusions from ECs act as a somatic-germline barrier that compartmentalizes germ cells to prevent undue influence from GSC niche signals. A similar hypothesis has also been made with regard to fly testes, wherein somatic cyst cells (the counterparts of ECs) wrap GSCs and their progeny to facilitate proper differentiation[52, 61]. This type of somatic permeability barrier for germ cell growth and differentiation is also probably conserved across species. In the developing mouse ovary, a group of pregranulosa cells, called biopotential pregranulosa cells (BPG), express Wnt4 and Wnt6; these cells later become granulosa cells that wrap germ cells [62], similar to *Drosophila* ECs. Although it is unclear if granulosa cells comprise a permeability barrier to compartmentalize germ cells and allow their differentiation, one study showed that hyperactivation of Wnt4/β-catenin signaling in the germline results in abnormal fetal development [63]. In mammalian testes, the early phase of spermatogenesis (GSCs and progenitor spermatogonia) occurs at the basal compartment of the seminiferous epithelium. This early phase is physically separated from the later phase of spermatogenesis, which occurs in the apical compartment, by an epithelial layer of somatic Sertoli cells called the blood-testicular barrier (the Sertoli cell barrier) [64, 65]. Because blood vessels, lymphatic vessels and nerves do not enter into the seminiferous epithelium, the blood-testicular barrier regulates the entry of molecules, such as nutrients and hormones, into the apical compartment in which germ cells enter meiosis [65]. Disruption of the blood-testicular barrier leads to a failure of spermatogenesis [65, 66].

A similar physical barrier of cells is found in the *C. elegans* gonad, but this barrier is formed by the germ cells themselves [67]. The germ cells in *C. elegans* form a syncytium in which the nuclei are enclosed by a partial plasma membrane, which has large openings on a central cytoplasmic core called “rachis”; some germ cells with partial membranes span the rachis and form cell bridges to pattern stemness Notch signaling in the gonad. Overall, these studies and ours strongly suggest that at least in the gonads, physical permeability barriers formed by cells can help to establish morphogen territories for proper cell patterning.

## Materials and Methods

### Fly strains and husbandry

Fly stocks were maintained at 22-25°C on standard medium, unless otherwise indicated. *y^1^w^1118^* was used as a wild-type control. The *bam^△86^* and *bam^1^* null alleles have been described previously [68]. *fz3RFP* (a gift from Dr. Rangan, Department of Biological Sciences University at Albany, State University of New York, USA) and *3GRH4TH-GFP (86FB)* (a gift from Dr. Cadigan, Department of Molecular, Cellular and Developmental Biology, University of Michigan, USA) were used to monitor Wnt signaling activity [32, 69]. Wg-GFP is an CRISPR/cas9-mediated in-frame insertion of GFP after the first exon of *wg* (a gift from Dr. Jean-Paul Vincent, The Francis Crick Institute, UK) [51]. Dpp-mcherry is a CRISPR/cas-9-mediated in-frame insertion of mCherry after amino acid 465 of Dpp (a gift from Dr. Thomas Kornberg, Cardiovascular Research Institute, UCSF, USA) [70]. *cycB3P-cycB3-gfp*, consisting of 6.5 Kb of the *cycB3* promoter driving the *cycB* coding region fused with GFP, was used to examine CycB3 expression (a gift from Dr. Dongsheng Chen, the Institute of Bioinformatics, College of Life Sciences, Anhui Normal University, China). *UAS-RNAi* lines against *tkv* (N#14026R-3 and V#3059), *wg* (B#32994 and V13352), *wnt2* (V#104338), *wnt4* (B#29442), *wnt5* (V#101621), *wnt6* (B#30493), *wnt8* (V#107727), *wnt10* (V#100867), *cycB3* (B#41979), *cycB* (B#34544), *cycE* (B#38920), *arm* (V#107344 (V1) and V#7767 (V2)), *dsh* (B#31306), *pygo* (V#100724), *axin* (B#62434), *daam* (V#24885), rhoA (B#28985), *rac1* (B#32383), and *gfp* (B#9331, second chromosome (II), and B#9330, third chromosome (III) were obtained from the National Institute of Genetics (N), Vienna Drosophila Resource Center (V), or Bloomington Drosophila Stock Center (B). The efficiencies of *RNAi* lines have been previously tested [11-13, 37, 41, 69, 71-76]. *UAS-arm^S10^* (B# 4782, a constitutively active form of Arm lacking a GSK3 phosphorylation site for degradation), was obtained from the Bloomington Drosophila Stock Center and has been described previously [12]. *UAS-arm-mGFP6* (B#58724) is the *arm* coding region linked via a polyserine linker to a C-terminal mGFP6 tag under the control of *UASp* regulatory sequences [77]. *bab1-GAL4*, *c587-GAL4*, *tj-GAL4* and *nos-GAL4* were used to drive transgene or RNAi expression; expression patterns of the somatic GAL4 drivers are summarized in Supplementary Table 1. Flies expressing *RNAi* driven by *tj-GAL4* for stage-specific experiments also carried *tub-GAL80^ts^* to control GAL4 expression; the flies were cultured at 18°C to silence GAL4 expression and were maintained at 29°C to allow GAL4 expression [39]. Other genetic tools are described in flybase (http://flybase.bio.indiana.edu).

### Developmental stage of larvae and pupae

The developmental stages of *Drosophila* were morphologically defined as previously described [78]. Flies were transferred to a new vial at 25°C to lay eggs for 3-6 h and then were removed. The vial was kept at 25°C. Newly hatched flies (First instar larvae, L1) were collected for dissection or further culturing. At approximately 50 h after egg laying (AEL), larvae were second-instar larvae (L2). Larvae climbing up and down from the food were considered mid-third instar larvae (ML3, 96 AEF), and larvae that had left the food and began wandering were late-third instar larvae (LL3, ∼120 AEL). Mid-and late-pupae were collected at around 170 and 194 AEL, respectively. Newly eclosed flies collected within 24 h were referred to as 1-day-old flies.

### Cloning and probe synthesis for *in situ* hybridization

Total RNA was extracted from 20 pairs of 1-day-old ovaries by using the GENEzol^TM^ TriRNA Pure Kit (Geneaid) according to the manufacturer’s manual. Total RNA (1 µg) was reversed transcribed with the Transcriptor First Strand cDNA Synthesis kit (Roche). Fragments of *fz3* and *wg* were amplified and used for the templates for synthesizing antisense probes; primers used are listed in the Supplementary Table 2. mRNA probes labeled with digoxigenin-UTP (Roche) were synthesized from 1 μg of the above PCR product using the ampliCap^TM^ SP6 high-yield message marker kit (Cell Script).

### Immunohistochemistry and fluorescence microscopy

For immunostaining, gonads and ovaries were dissected, fixed and immunostained at designated stages, as described previously [20]. In brief, ovaries were dissected in Grace’s insect medium (GIM) (Lonza) and fixed with 5.3% paraformaldehyde (PFA)/GIM for 13 min, Then, samples were washed in PBST (0.1% Triton X-100 in PBS) three times for 20 min each and teased apart in PBST. Samples were incubated in blocking solution (GOAL Bio) for 3 h at room temperature (RT) or 4°C overnight (O/N). Samples were then incubated with primary antibodies (diluted in blocking solution) for 5 h at RT or 4°C overnight, followed by four PBST washes for 30 min each. Samples were incubated with secondary antibodies (diluted in blocking solution) for 5 h at RT or 4°C O/N, followed by four PBST washes for 30 min each. Primary antibodies were used as follows: mouse anti-Hts (Drosophila Studies Hybridoma bank, DSHB, 1B1, 1:50), mouse anti-α-Spectrin (DSHB 3A9, 1:100), mouse anti-Lamin (Lam) C (DSHB LC28.26, 1:25), guinea pig anti-Traffic Jam (1:10000; a gift from Dorothea Godt, University of Toronto, Canada), rabbit anti-Vasa (Santa Cruz Sc-30210, 1:500), and chicken anti-GFP (Invitrogen, A10262, 1:1000). Secondary antibodies were used as follows: Alexa Flour 488 anti-rabbit IgG (Invitrogen, 1:250), Alexa Flour 488 anti-mouse (Invitrogen, 1:500), Alexa Flour 568 anti-mouse IgG (Invitrogen, 1:250), Alexa Fluor 647 anti-anti-Guinea Pig IgG (Invitrogen, 1:250), Alexa Flour 488 anti-chicken (Jackson, 1:1000), DNA was stained with 0.5 μg/ml DAPI (Sigma) or TO-PRO-3 (Invitrogen) for 10 min at RT or O/N at 4°C. Finally, samples were mounted in 80% glycerol containing 20 µg/ml N-propyl gallate (Sigma) or Vectashield mounting medium (Vector Laboratories) and analyzed using a Zeiss LSM 700, 900 or Leica SP8 confocal microscope.

Fluorescent RNA *in situ* hybridization was performed as described in a previous report [79], with slight modifications. In brief, larval or adult ovaries were dissected in GIM and fixed by 4% PFA in PBS (DEPC-treated) with freshly added 1% DMSO for 20 min at RT or O/N at 4°C. Samples were washed in PBS and dehydrated through a series of ethanol solutions (25%, 50%, 75%, and 100%), followed by storage at −20°C. Samples were rehydrated through a series of ethanol solutions (100%, 75%, 50%, and 25%), rinsed by PBS, and treated with 50 μg/ml Proteinase K for 5 min. After a post-fixation step in 4% PFA in PBST (1X PBS with 0.1% Tween 20) for 30 min at RT, samples were washed well and prehybridized in hybridization buffer (HYB^+^) (50% formamide, 5X SSC, 50 μg/ml heparin, 0.1% Tween-20, 100 μg/ml tRNA, 10 μg/ml Salmon Sperm DNA) for 1 h at 60°C. Then, samples were hybridized in HYB^+^ containing denatured DIG-labeled RNA probes (100-150 ng) at 60°C O/N. Samples were washed with a series of HYB^-^ (50% formamide, 5X SSC with 0.1% Tween) mixed into 2X SSC (0.3M NaCl, 30mM sodium citrate) (75%, 50%, and 25%) at 65°C and a series of 0.2X SSC solutions (75%, 50%, and 25%) at 68°C, followed by a rinse with PBST at RT. Samples were treated with 3% H_2_O_2_/PBT for 1 h at RT to inactivate endogenous perioxidase (POD), and then the samples were blocked in 2X Roche Blocking solution for 1 h at RT. Ovaries were incubated with anti-Dig-POD (1:500, Roche # 11207733910) in blocking buffer at 4°C O/N, washed well, and incubated in 1:200 TSA/amplification buffer (TSA Plus Fluorescence Kits; PerkinElmer) for 30 min to develop signals. After washing, ovaries were blocked with blocking solution (GOAL Bio), and then the immunostaining procedure was followed, as described above.

### Imaging quantification

GSCs were defined by their location directly adjacent to niche cap cells, and their fusome that is juxtaposed to the GSC-cap cell junction. SCCs were identified as cells with round-shaped fusomes (CBs in controls) that are not GSCs. To measure *fz3-RFP* expression, five confocal z-sections of each germarium carrying a clear EC region (5 sections) were merged and analyzed with ImageJ. The EC region was selected and the mean intensity (arbitrary units) was measured. To assess CycB3-GFP expression, Zen 3.1 (blue edition, ZEISS) was used to measure the mean fluorescence intensity of the confocal z-section with the largest nuclear diameter from each GFP-positive germ cell. To measure *fz3* transcript signals in the germ cell, 4-5 confocal z-sections covering the largest area of the germarium were assessed. The numbers of signals in germ cells marked by Vasa-GFP were counted using ImageJ. To measure GFP-Wg signals, confocal z-sections containing images of TF and cap cells or the germ cell region were merged, and numbers of GFP-Wg granules were counted using ImageJ.

Each experiment was performed with at least two biological replicates. For fixed samples, 10 newly eclosed females with the indicated genotypes were randomly picked from a standard cross; ovaries were dissected and subjected to immunostaining. Ovaries from at least 10 pairs of ovaries were separated, mixed, and mounted for observation; 5-15 images were collected of representative phenotypes for each replicate. Statistical analysis was performed as described in the figure legend.

### Live imaging of adult germaria

Live images were captured of germaria carrying Wg-GFP, Dpp-mcherry or cycB3p-CycB3-GFP, as fixation caused high background that interfered with analysis of signals. To obtain images of live adult germaria, ovaries of 1-day-old flies were dissected in GIM, and the anterior portions of the ovarioles were gently teased apart to separate germaria. Ovaries were then stained with or without CellMask™ Deep Red Plasma Membrane Stain (1:2000 diluted in GIM, Invitrogen, C10046) for 1 min at RT. A short incubation time was used to prevent/reduce the staining of germ cell membranes. Note that somatic *tkv*-KD and *bam* mutant germaria have blunted EC protrusions, and the CellMask signal within those germaria likely corresponds to germ cell membranes. Ovaries were transferred on to a glass slide, and 15 µl fresh GIM was added; sheath cells were removed from each ovariole using a tungsten filament needle. For the permeability assay, after removing the sheath, 0.3 µl of 5 µg/µl 10-kDa dextran conjugated with Alexa Fluor™ 488 (Invitrogen, D22910, a gift provided by Dr. Y. Henry Sun, Institute of Molecular Biology, Academia Sinica, Taiwan) was directly added to the 15 µl GIM on the slide (final concentration, 0.2 µg/µl). Finally, the ovarioles were covered with a coverslip and imaged using a Zeiss LSM 900 confocal microscope.

### Chromatin immunoprecipitation (ChIP) assay

The ChIP assay was performed as previously described, with minor modifications [20]. In brief, 100 pairs of 7-day-old *nos>gfp* and *nos>arm-mGFP6* ovaries of flies kept at 29°C were dissected in cold GIM. The ovaries were fixed in 950 μl PBS containing 1.8% formaldehyde for 10 min at RT. Cross-linking was stopped by adding 50 μl of 2.5 M Glycine. Fixed ovaries were ground in cold Buffer A1 (15 mM Hepes, pH 7.5, 15 mM NaCl, 60 mM KCl, 4 mM MgCl_2_, 0.5% Triton X-100, 0.5 mM DTT, 1 mM PMSF, 5 mM NaF, Protease inhibitor). Chromatin pellets were precipitated by centrifugation at 1800 ×*g* for 5 min at 4°C and washed with buffer A1 three times. Chromatin pellets were then washed one time with buffer A2 (15 mM Hepes, pH 7.5, 140 mM NaCl, 1 mM EDTA, 0.5 mM EGTA, 1% Triton X-100, 0.1% sodium deoxycholate, 0.1% SDS, 0.5% N-lauroyl sarcosine, 1 mM PMSF, 5 mM NaF and 1x Protease inhibitor). Chromatin pellets were then sonicated in 450 μl buffer A2 using a Bioruptor (Diagenode) for 16 min (30 s on/30 s off). Chromatin solutions were obtained by centrifugation, 14,000 rpm, 10 min, at 4°C after sonication. Twenty-five microliters of GFP-Trap bead slurry (GFP-Trap Magnetic Agarose, Chromtek, gtma-10) was added to 500 μl chromatin solution and incubated overnight at 4°C. GFP-Trap beads were washed with 500 μl RIPA buffer (10 mM Tris-HCl, pH 7.5, 150 mM NaCl, 0.5 mM EDTA, 1% Triton X-100, 0.1% SDS, 1% sodium deoxycholate) three times and twice with TE buffer. The chromatin was eluted twice in TE buffer containing 1% SDS and 250 mM NaCl for 20 min at 65°C. Eluted chromatin solutions were treated with RNase A and proteinase K, then cross-linking was reversed overnight at 65°C. DNA was purified using a QIAquick PCR Purification Kit (QIAGEN). Input and immunoprecipitated DNA samples were used for qRT-PCR with qPCRBIO SyGreen Mix (PCR Biosystems). The primers used to amplify fragments of the *cycB3* and *rp49* gene are listed in Supplementary Table 1.

### Western blot analysis

Forty pairs of ovaries were dissected from newly enclosed flies with little or no stage 14 egg chambers. Samples were lyzed and homogenized on ice in RIPA buffer (20 mM Tris HCl, pH 7.5, 150 mM NaCl, 1% NP-40, 1mM EDTA) supplemented with 2× EDTA-free Complete Protease Inhibitor Cocktail (Roche). Protein lysates were collected from supernatant after centrifugation at 4 °C, 12000 rpm for 5 min. Lysate was added into same volume 2× Laemmli sample buffer (126 mM Tris/Cl, pH 6.8, 20% glycerol, 4% SDS and 0.02% bromophenol blue) containing 10% β-mercaptoethanol and then boiled for 10 min at 70 °C in, separated by 10% SDS polyacrylamide gels (SDS-PAGE) and blotted onto polyvinylidene difluoride (PVDF) membranes. Membranes were blocked by 5% skim milk in 1× Tris-buffered saline containing 0.1% Triton X-100 (TBST, pH 7.5) for 1 h at room temperature, then incubated with rabbit anti-GFP (1:2000, Torrey Pines Biolabs, #TP401), mouse anti-Wg 4D4 (1:2000, DSHB) in TBST containing 1% skim milk at 4 °C overnight with shaking. After three 10 min washes with 1× TBST, membranes were incubated with horseradish peroxidase (HRP)-conjugated goat anti-rabbit IgG (1:5000, Jackson ImmunoResearch), HRP-conjugated goat-anti-mouse IgG (1:10000, Croyez Bioscience Co., Ltd.) in TBST containing 1% skim milk for 1 h at room temperature. After three 10 min washes with 1× TBST, signals were detected by chemiluminescence with a Western LightningTM Plus-ECL kit (PerkinElmer).

## Acknowledgements

We thank D. Godt, P. Rangan, K. Cadigan, H. Sun, JP Vincentthe, DS Chen, T. Kornberg, Bloomington, National Institute of Genetics-Fly Stocks, and VDRC Stock Center, and the DSHB for *Drosophila* stocks and antibodies. We also thank M. Calkins for English editing. This work was supported by and the Ministry of Science and Technology, Taiwan (107-2311-B-001-004-MY3), and the intramural funding from the Institute of Cellular and Organismic Biology, Academia Sinica, Taiwan.

## Author Contributions

SMY, KYL, TAC, CYT, CHL, YTW, LL, YC and HJH conceived and designed the experiments. SMY (Fig. 1-3, 6 and 7, and Supplementary Fig. 1-4), KYL (Fig. 1, 5-7, Supplementary Fig. 5-8), and TAC (Fig. 4-6, and Supplementary Fig. 2, 6-9), YTW (Fig. 3 and 4), CYT (Fig. 4 and 7), CHL (Fig. 5K-M) and LL (Supplementary Fig. 10) performed the experiments. SMY, KYL, YC and HJH analyzed the data and wrote the manuscript. The authors declare no competing financial interests.

