## Supplemental figures for "Canonical Wnt signaling promotes formation of somatic permeability barrier for proper germ cell differentiation"

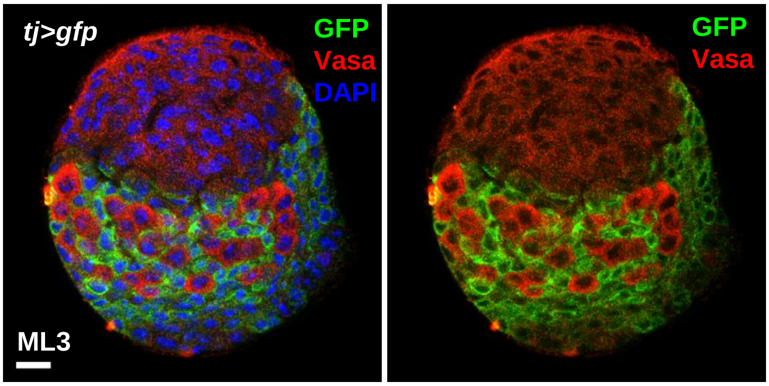


**Supplementary Fig. 1**

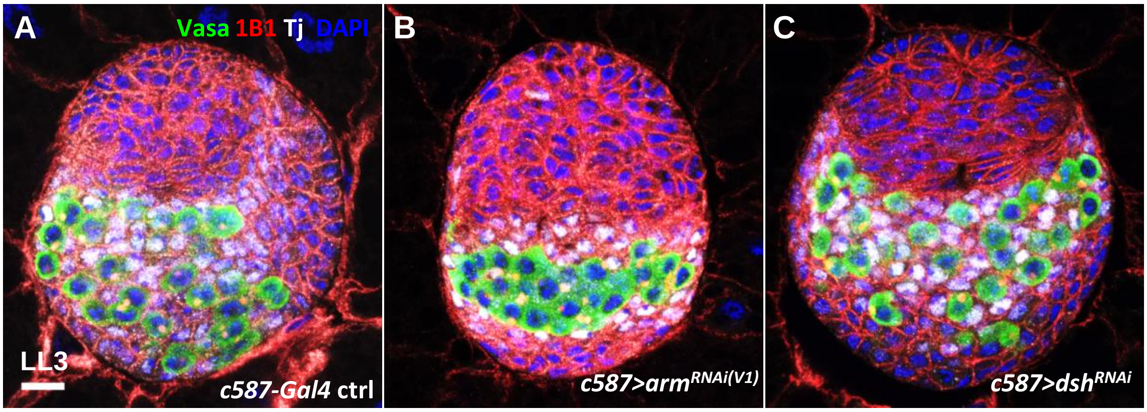


**Supplementary Fig. 2**

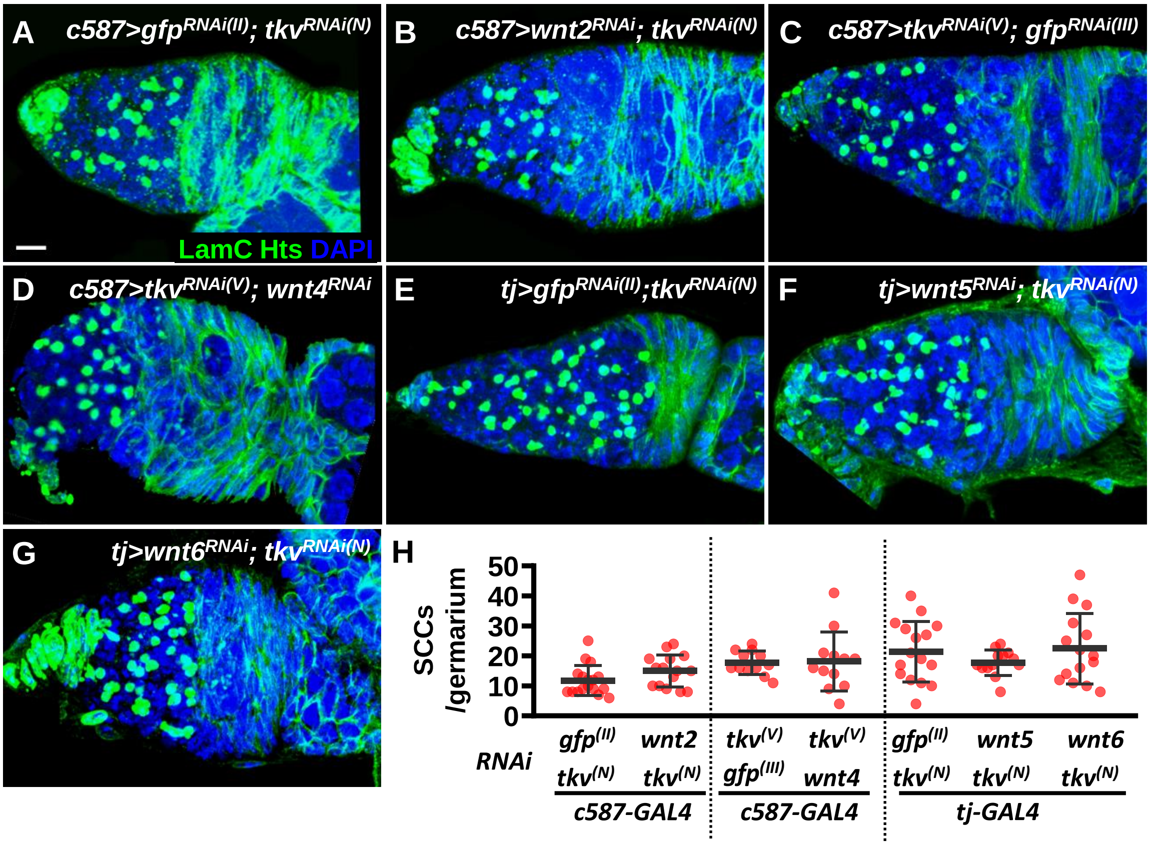


**Supplementary Fig. 3**

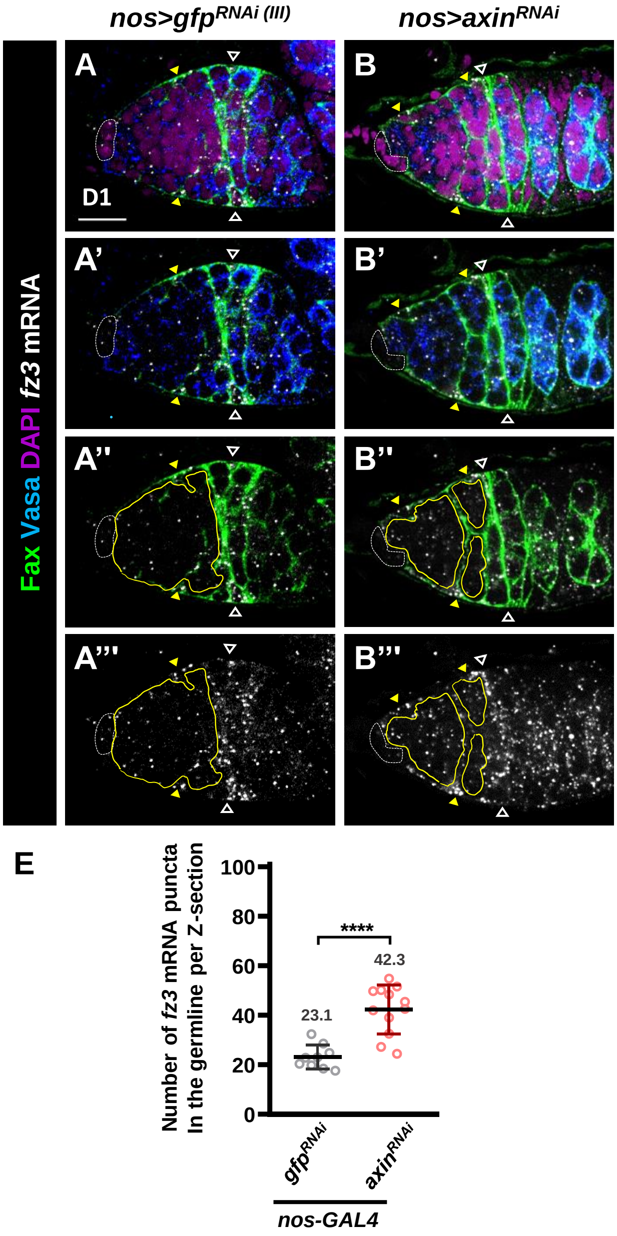


**Supplementary Fig. 4**

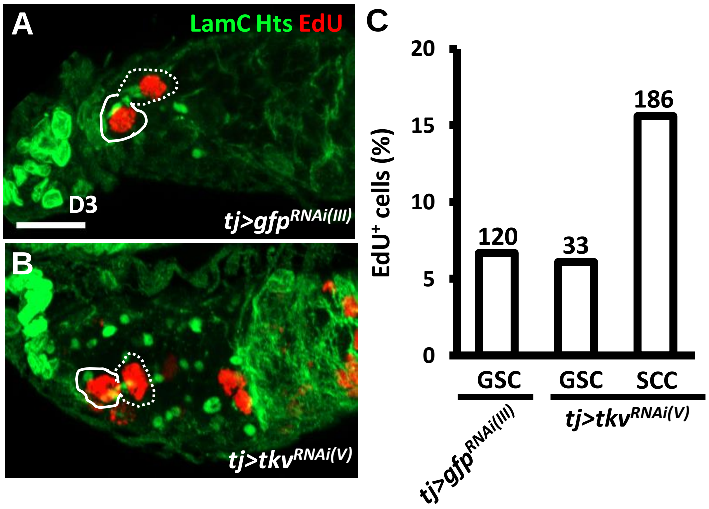


**Supplementary Fig. 5**

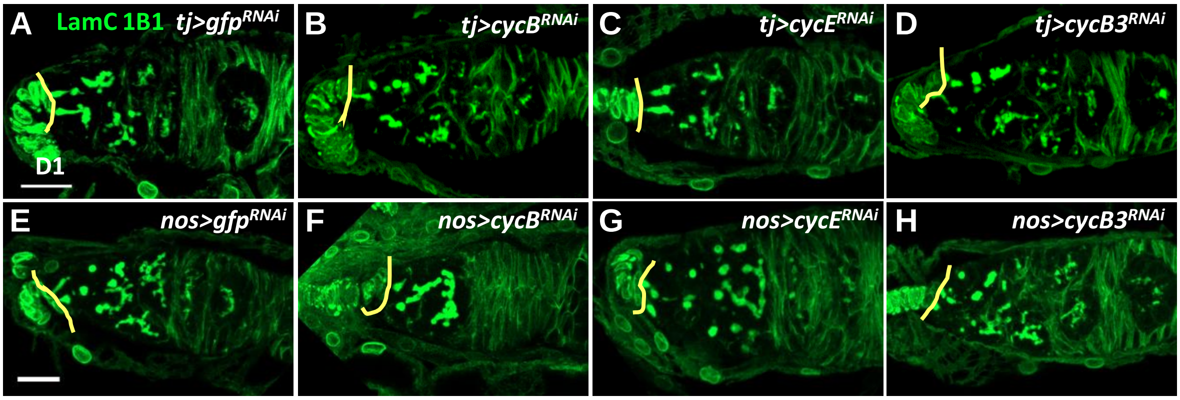


**Supplementary Fig. 6**

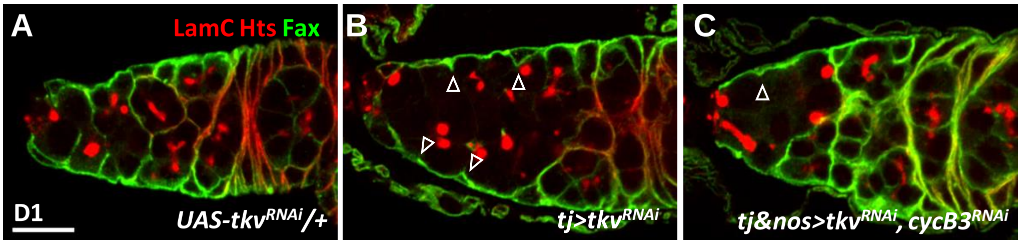


**Supplementary Fig. 7**

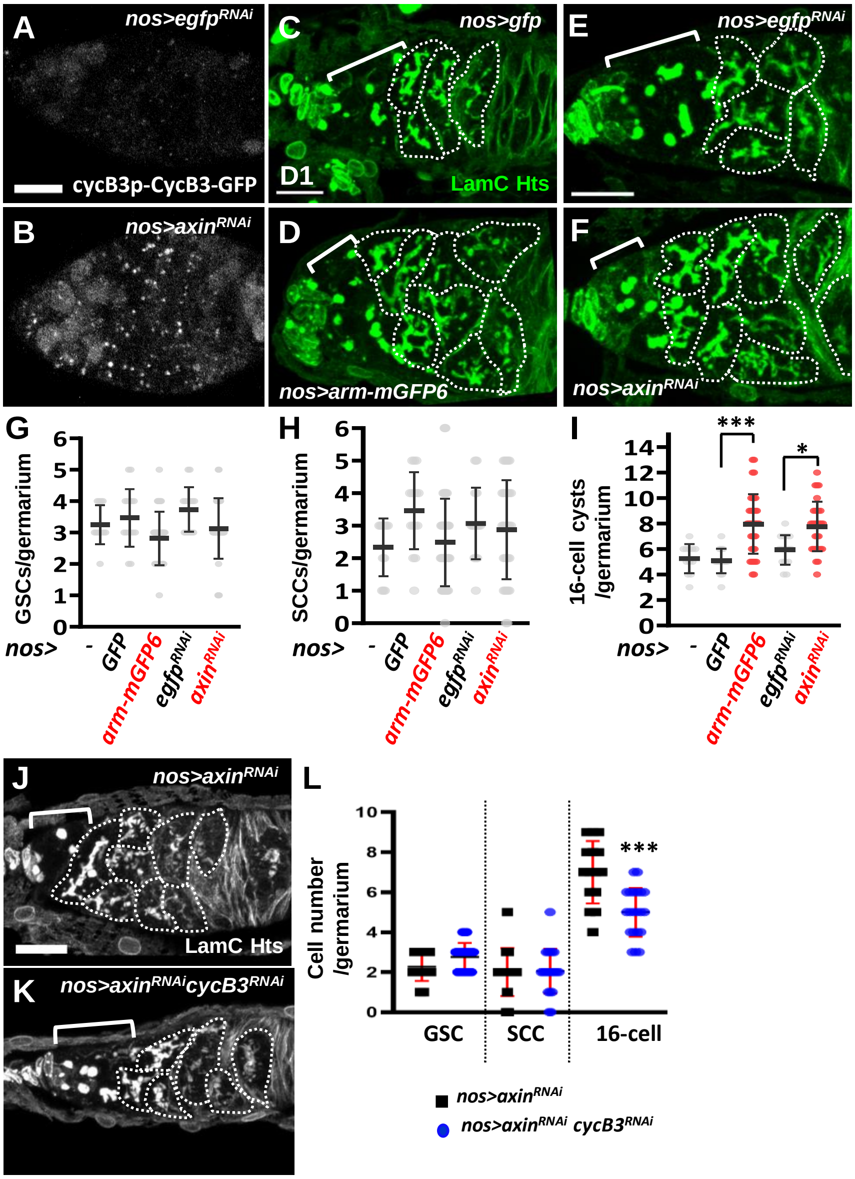


Supplementary Fig. 8

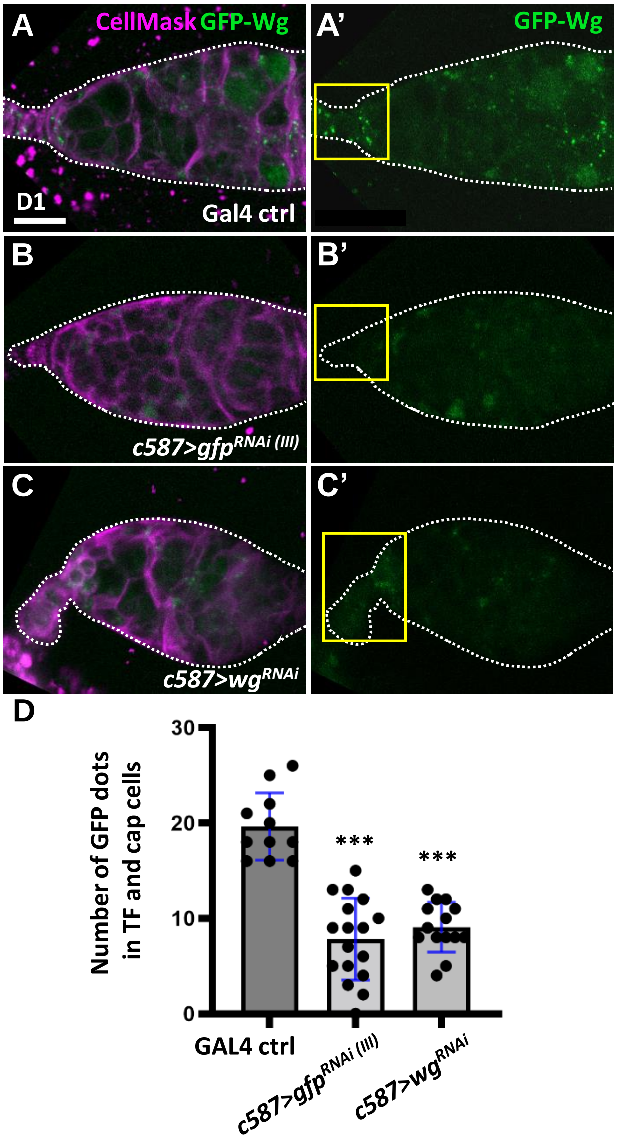


**Supplementary Fig. 9**

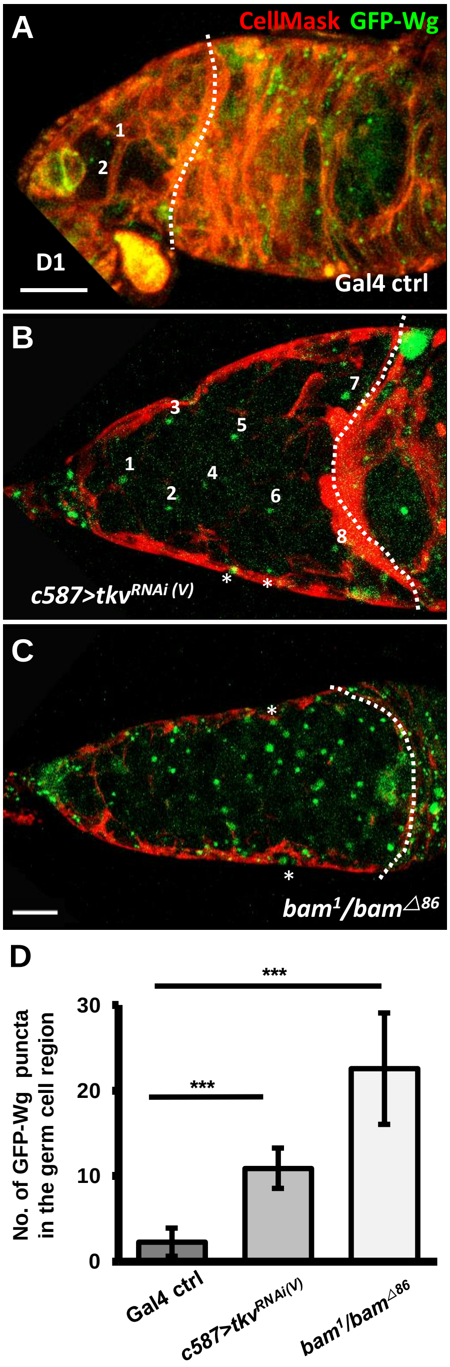


**Supplementary Fig. 10**

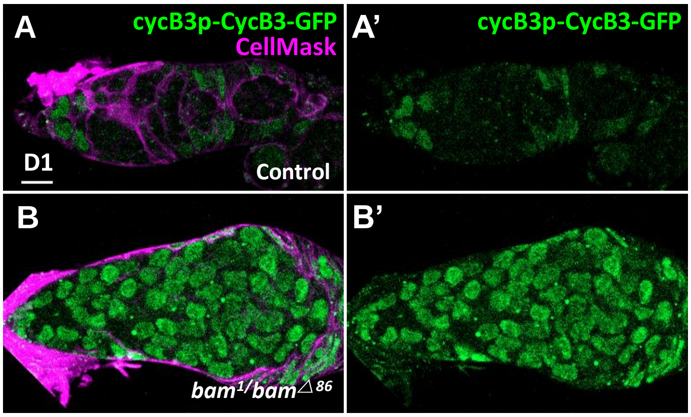


**Supplementary Fig. 11**

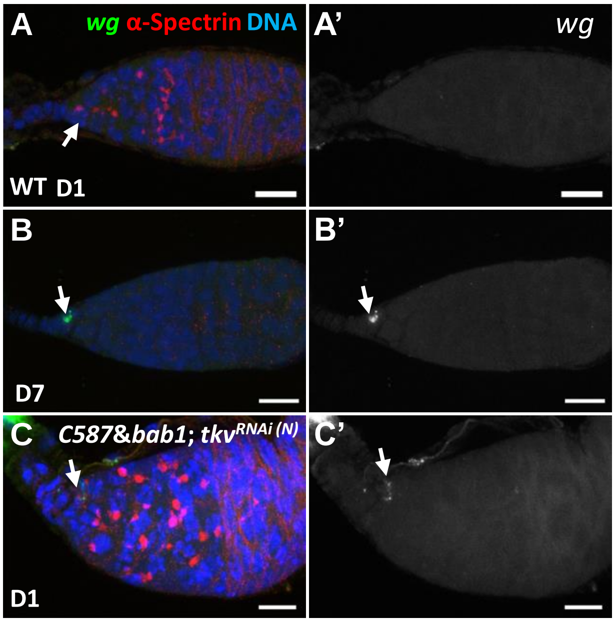


**Supplementary Fig. 12**

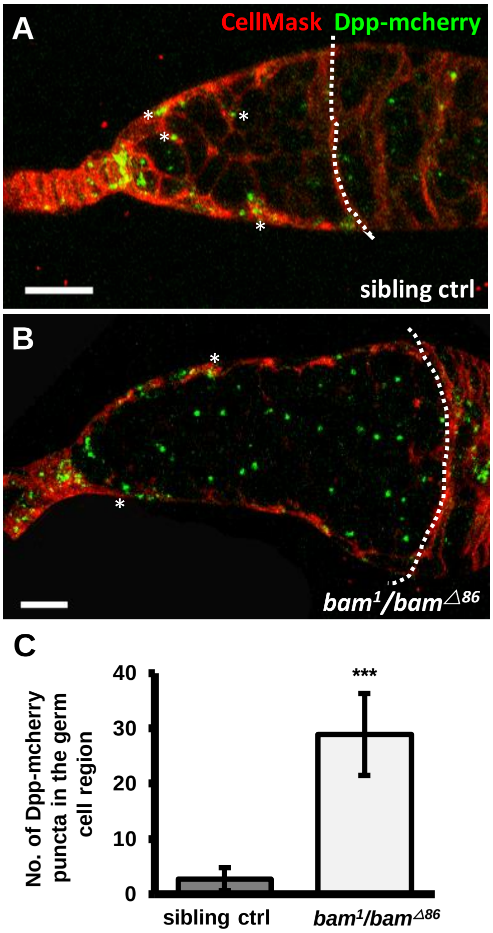


**Supplementary Fig. 13**

### Supplementary Figure legends

**Supplementary. Fig. 1 *tj-GAL4* is activated in intermingled cells of the developing ovary.** Mid-L3 (ML3) ovary expressing GFP (green) driven by *tj-Gal4* with Vasa (red, germ cells) and DAPI (blue, DNA). GFP was expressed throughout development until dissection. Scale bar, 10 µm.

**Supplementary Fig. 2 Somatic knockdown of *arm* disrupts soma-germline interaction. (A-C)** Late-L3 (LL3) c*587-Gal4* control (ctrl) (A), c587>$arm^{RNAi(V1)}$ (B) and c587>$dsh^{RNAi}$ (C) ovary with Vasa (green, germ cells), Hts (red, fusomes), Tj (gray, intermingled cells), DAPI (blue, DNA). *RNAi* was expressed throughout development until dissection. Genotype of the control in A is *c587-Gal4/+*, Scale bar, 10 µm.

**Supplementary Fig. 3 Somatic co-knockdown of *tkv* with *wnt2, wnt4, wnt5* or *wnt6* does not rescue germ cell differentiation defect. (A-G)** One-day (D)-old *c587>*$gfp^{RNAi(II)}$ & $tkv^{RNAi(N)}$ (A), *c587>*$wnt2^{RNAi}$ & $tkv^{RNAi(N)}$ (B), *c587>*$tkv^{RNAi(V)}$ & $gfp^{RNAi(III)}$ (C), *c587>*$tkv^{RNAi(V)}$ & $wnt4^{RNAi}$ (D), *tj>*$gfp^{RNAi(II)}$ & $tkv^{RNAi(N)}$ (E), *tj>*$wnt^{RNAi}$ & $tkv^{RNAi(N)}$ (F), and *tj>*$wnt6^{RNAi}$ & $tkv^{RNAi(N)}$ germaria (G) with LamC (green, terminal filament and cap cell nuclear envelopes), Hts (green, fusomes) and DAPI (blue, DNA). Scale bar, 10 µm. **(H)** Number of spectrosome-containing cells (SCCs) in the indicated genotypes. Error bar, mean ± SD. Statistical analysis, One-way ANOVA. *RNAi* was expressed throughout development until dissection.

**Supplementary Fig. 4 *fz3* transcripts are increased in the germline upon *axn* knockdown.** **(A and B)** *In situ* hybridized *tj>gfp*$^{RNAi\ (III)}$ (A) and *tj>axin*$^{RNAi}$ germaria (B) with Fax (green, escort cell membrane extension), Vasa-GFP (blue, germ cells), and *fz3* mRNA (gray) and DAPI (magenta, DNA). A' and B' show Fax3 and Vasa-GFP signal channels; A'' and B'' show Fax3 and *fz3* mRNA channels; A''' and B''' show *fz3* mRNA channel. Hollow triangles point to the 2A/B boundary; yellow triangles indicate escort cell region; germ cell region before the 2A/B boundary are outlined by yellow circles in A'', A''', B'' and B'''. **(O)** Number (No.) of *fz3* mRNA puncta in the germline per germarium with indicated genotype. *RNAi* was expressed throughout development. Scale bar, 10 µm. *, $P < 0.05$, **, $P < 0.01$; ***, $P < 0.001$. Error bars indicate mean ± S.D., Student's *t* test was used for statistical analysis.

**Supplementary Fig. 5 Spectrosome-containing cells exhibit proliferation capacity in the soma *tkv*-KD germaria. (A and B)** One-day-old *tj>gfp*$^{RNAi}$ (A) and *tj>tkv*$^{RNAi(V)}$ germaria (B) with Hts (green, fusomes), LamC (green, terminal filament and cap cell nuclear envelopes), and EdU (red, S phase marker). Scale bar, 10 µm. **(C)** Percentage (%) of EdU-positive (+) GSCs or SCCs in the indicated genotypes. 55 control and 9 *tj>tkv*$^{RNAi}$ germaria were analyzed; number of cells analyzed is shown above each bar.

**Supplementary Fig. 6 Germline and somatic knockdown of *cycB*, *cycB3* and *cycE* throughout development. (A-D)** One-day (D)-old *tj>gfp*$^{RNAi(III)}$ (A), *tj>cycB*$^{RNAi}$ (B), *tj>cycE*$^{RNAi}$ (C) and *tj>cycB3*$^{RNAi}$ germaria (D) do not show obvious phenotypes. **(E-H)** One-day (D)-old *nos>gfp*$^{RNAi(III)}$ (E), *nos>cycB*$^{RNAi}$ (F), *nos>cycE*$^{RNAi}$ (G) and *nos>cycB3*$^{RNAi}$ germaria (H). Germline knockdown of *cycB* reduced the number of GSCs, while germline knockdown of cycE caused GSC loss and differentiation defect. Germaria labelled with Hts (green, fusomes), LamC (green, terminal filament and cap cell nuclear envelopes). Solid lines mark the junction between cap cells and GSCs. Scale bar, 10 μm.

**Supplementary Fig. 7 Knockdown of *cycB3* in the germline of somatic-*tkv* knockdown germaria partially rescues germ cell differentiation and escort cell protrusions. (A-C)** One-day (D)-old *UAS-tkv*$^{RNAi(V)}$*/+* (A), *tj>tkv*$^{RNAi\ (V)}$ (B), *tj & nos>tkv*$^{RNAi\ (V)}$ *& cycB3*$^{RNAi}$ (C) with Hts (red, fusomes and follicle cell membranes) and Fax (green, escort cell protrusions). Arrowheads point to blunted EC protrusions. Scale bar, 10 μm.

**Supplementary Fig. 8 Hyperactivation of germline Wnt signaling promotes germ cell differentiation via CycB3. (A and B)** Live images of *nos>gfp*$^{RNAi(III)}$ (A) and *nos>axin*$^{RNAi}$ (B) with cycB3P-CycB3-GFP (gray). **(C-F)** *nos>GFP* (C), *nos>Arm-mGFP6* (D), *nos>gfp*$^{RNAi(III)}$ (E), *nos>axn*$^{RNAi}$ (F) with LamC (green, terminal filament and cap cell nuclear envelopes), Hts (green, fusomes). **(G-I)** Numbers of GSCs (G), SCCs (H) and 16-cell cysts (I) in the indicated genotypes.

**(J and K)** *nos>axin*$^{RNAi}$ (J) and *nos>axin*$^{RNAi}$*cycB3*$^{RNAi}$ (K) with LamC (gray) and Hts (gray). (L) Numbers of GSCs, SCCS, and 16-cell cysts per germarium in the indicated genotypes. Brackets indicate region 1 of the germarium; 16-cell cysts are marked by dashed circles. Scale bar, 10 μm. Statistical analysis in G-I was by one-way ANOVA, in L Student's *t*-test. **P* < 0.05; ****P* < 0.001.

**Supplementary Fig. 9 GFP-Wg expression mimics endogenous Wg expression. (A-C)** One-day (D)-old *c587-GAL4/+* (A), *c587>gfp*$^{RNAi\ (III)}$ (B), *c587>wg*$^{RNAi}$ germaria (C) bearing GFP-Wg (green) with CellMask (magenta, cell membrane). A'-C' only show GFP-Wg channel. Scale bar, 10 μm. **(D)** Number of GFP-Wg granules in the TF and cap cell region of control, *c587>gfp*$^{RNAi\ (III)}$, and *c587>wg*$^{RNAi}$ germaria. ***, *P*<0.001. Statistical analysis was by Student's *t*-test.

**Supplementary Fig. 10 More GFP-Wg granules are present in the germ cell zone of germaria with blunted escort cell protrusions. (A-C)** Live images of one-day (D)-old GAL4 control (A), c587>*tkv*$^{RNAi(V)}$ (B), and *bam1/bam*$^{\triangle 86}$ germaria (C) merged from 8 z-sections (9 micron). Germaria expressing GFP-Wg (green) labeled with CellMask (red, membrane dye). Arabic numerals reveal the GFP-Wg granules shown in the germ cell region before 2A/2B boundary (A), or the junction between escort cells and follicle cells (B). GFP-Wg granules in the bam$^{1}$/bam$^{\triangle 86}$ germarium are not shown. Asterisks mark GFP-Wg in ECs. **(D)** Number (no) of GFP-Wg puncta in the germ cell region before the 2A/2B boundary of control germaria, or before the junction between escort cells and follicle cells of c587>*tkv*$^{RNAi(V)}$, and *bam1/bam*$^{\triangle 86}$ germaria. Scale bars, 10 μm. ***, *P*<0.001. Statistical analysis was by Student's *t*-test. The genotype of GAL4 control is *c587>gfp*$^{RNAi(III)}$*, GFP-wg/+*.

**Supplementary Fig. 11 CycB3-GFP expression is present in SCCs of *bam* mutant germaria. (A and B)** Live images of 1-day (D)-old control (A) and *bam*$^{1}$*/bam*$^{\triangle 86}$ germaria (B) bearing *cycB3P-cycB3-gfp* (Green, CycB3-GFP) with CellMask (magenta, cell membrane). Scale bar, 10 μm. The genotype of the control is *cycB3p-cycB3-gfp/+*.

**Supplementary Fig. 12 *wg* expression pattern is not altered in somatic *tkv*-KD germaria. (A -C)** control (A and B) and somatic-*tkv* KD germaria (C) with *wg* RNA (green), α-Spectrin (red, fusomes) and TO-PRO-3 (blue, DNA). The genotype in A and B is *yw*, and in C is *c587-GAL4/+; bab1-GAL4/UAS-wg*$^{RNAi}$. Wg is nearly undetectable in newly eclosed (D1) control flies and can be clearly detected in cap cells of 7-day-old germaria. Two somatic GAL4 drivers were simultaneously used to enhance *tkv* knockdown in the soma throughout development. It is not clear why *tkv* knockdown increases *wg* transcripts in cap cells. A and C are one-day (D)-germaria; B is 7-day-old germarium. Arrows indicate cap cells. Scale bar, 10 μm.

**Supplementary Fig. 13 More Dpp-mcherry granules are present in the germ cell zone of germaria with blunted escort cell protrusions. (A and B)** Live images of one-day (D)-old sibling control (A) and *bam1/bam*$^{\triangle 86}$ germaria (B) merged from 18 z-sections (12 micron). Germaria expressing Dpp-mcherry (green) labeled with CellMask (red, membrane dye). Asterisks mark GFP-Wg in ECs. **(C)** Number (no) of Dpp-mcherry puncta in the germ cell region before the 2A/2B boundary of control germaria, or before the junction between escort cells and follicle cells of *bam1/bam*$^{\triangle 86}$ germaria. ***, $P$<0.001. Statistical analysis was by Student's *t*-test. Scale bars, 10 μm.
